# ASAR lncRNAs control DNA replication timing through interactions with multiple hnRNP/RNA binding proteins

**DOI:** 10.1101/2022.06.04.494840

**Authors:** Mathew J. Thayer, Michael B. Heskett, Leslie G. Smith, Paul T. Spellman, Phillip A. Yates

## Abstract

ASARs are a family of very-long noncoding RNAs that control replication timing on individual human autosomes, and are essential for chromosome stability. The eight known ASAR lncRNAs remain closely associated with their parent chromosomes. Analysis of RNA-protein interaction data (from ENCODE) revealed numerous RBPs with significant interactions with multiple ASAR lncRNAs, with several hnRNPs as abundant interactors. An ∼7kb domain within the ASAR6-141 lncRNA shows a striking density of RBP interaction sites. Genetic deletion and ectopic integration assays indicate that this ∼7kb RNA binding protein domain contains functional sequences for controlling replication timing of entire chromosomes in *cis*. shRNA-mediated depletion of 10 different RNA binding proteins, including HNRNPA1, HNRNPC, HNRNPL, HNRNPM, HNRNPU, or HNRNPUL1, results in dissociation of ASAR lncRNAs from their chromosome territories, and disrupts the synchronous replication that occurs on all autosome pairs, recapitulating the effect of individual ASAR knockouts on a genome-wide scale. Our results further demonstrate the role that ASARs play during the temporal order of genome-wide replication, and we propose that ASARs function as essential RNA scaffolds for the assembly of hnRNP complexes that help maintain the structural integrity of each mammalian chromosome.

## Introduction

The vast majority of mammalian DNA replicates in homologous regions of chromosome pairs in a highly synchronous manner ^1–3^. However, genetic disruption of non-protein coding ASAR (“<u>AS</u>ynchronous replication and <u>A</u>utosomal <u>R</u>NA”) loci causes a delay in replication timing on individual human chromosomes in *cis*, resulting in highly asynchronous replication between pairs of autosomes ^4–7^. There are eight genetically validated ASARs that are located on human chromosomes 1, 6, 8, 9 and 15 ^4–7^. All of the known ASAR lncRNAs share several distinctive characteristics, including: 1) contiguously transcribed regions of >180 kb, 2) epigenetically regulated <u>A</u>llelic <u>E</u>xpression Imbalance (AEI; including mono-allelic expression); 3) <u>V</u>ariable <u>E</u>pigenetic <u>R</u>eplication <u>T</u>iming (VERT; including asynchronous replication timing); 4) high density of LINE1 (L1) sequences (>30%); and 5) retention of the RNA within their parent chromosome territory. Recently, we used these notable characteristics to identify 68 ASAR candidates on every human autosome ^5,6^.

In addition to genetic disruption assays, ectopic integration of ASAR transgenes into mouse chromosomes has been used as a second functional assay to help define the critical sequences within ASAR lncRNAs that control chromosome-wide replication timing ^8,9^. For example, ASAR6 lncRNA expressed from transgenes, ranging in size from an ∼180 kb BAC transgene to PCR products (as small as ∼3 kb) containing the critical sequences, remain associated with the chromosome territories where they are transcribed ^8,9^. These same transgenes cause <u>d</u>elayed replication timing (DRT) and <u>d</u>elayed <u>m</u>itotic <u>c</u>ondensation (DMC) of entire chromosomes in *cis* ^8,9^. Furthermore, ASAR6 lncRNA was found to control chromosome-wide replication timing using oligonucleotide mediated RNA degradation of ASAR6 lncRNA that was expressed from a BAC transgene integrated into mouse chromosome 3 ^9^. These findings suggest that ASARs represent ubiquitous, essential, *cis*-acting elements that control mammalian chromosome replication timing via expression of chromosome associated RNAs. In this report, we identify <u>R</u>NA <u>b</u>inding <u>p</u>roteins (RBPs) that are critical for ASAR lncRNA localization and function.

Chromosome-associated long non-coding RNAs are known to be important for numerous aspects of chromosome dynamics in diverse eukaryotes from yeast to mammals ^10–14^. The mammalian nucleus is rich in heterogenous nuclear RNA (hnRNA), which represents >95% of the total RNA Polymerase II output ^12,15^. The vast majority of hnRNA is poorly defined, has been referred to as the “Dark Matter” of the genome ^15^, and includes spliced and unspliced intronic sequences, circular RNAs ^16^, <u>v</u>ery long intergenic <u>n</u>on-<u>c</u>oding RNAs (vlincRNAs; ^17^), as well as poorly defined RNAs that include <u>c</u>hromatin <u>a</u>ssociated RNAs (caRNAs; ^18^) and <u>rep</u>eat rich RNAs (repRNAs ^19^) that can be detected by C0T-1 DNA (which is comprised primarily of LINE and SINE repetitive sequences; ^13,20,21^). Some of these nuclear RNAs comprise abundant species, yet lack clearly defined functions, while others are thought to be the by-products of other nuclear processes and have been collectively called “RNA debris” ^12^. The relationship between ASAR lncRNAs and these other chromosome-associated RNAs remains poorly defined.

Previous studies have suggested that long nascent transcripts that are detected by C0T-1 DNA play a dynamic structural role that promotes the open architecture of active chromosome territories ^13,20^. The RNA species detected by C0T-1 DNA are predominantly L1 sequences, and these L1 RNAs remain associated with the chromosome territories where they are transcribed ^20^. A link between C0T-1 RNA and ASAR lncRNAs is suggested by the observation that ASAR lncRNAs contain a high L1 content and remain associated with their parent chromosome territories ^4–7,9^. In addition, deletion and ectopic integration assays demonstrated that the critical sequences for controlling chromosome-wide replication timing within ASAR6 and ASAR15 map to the antisense strand of L1 sequences located within the ASAR6 and ASAR15 lncRNAs ^9^. In this report we found that human C0T-1 DNA can detect the chromosome territory localization of ASAR6 lncRNA that is expressed from a BAC transgene integrated into a mouse chromosome, indicating that at least some of the RNA species detected by C0T-1 DNA represents ASAR lncRNA.

Heterogeneous nuclear ribonucleoproteins (hnRNPs) represent a large family of abundant RBPs that contribute to multiple aspects of nucleic acid metabolism. Many of the proteins that are in hnRNP complexes share general features, but differ in domain composition and functional properties (reviewed in ^22^). The functions of hnRNPs vary according to their subcellular localization. Most of the hnRNPs possess conventional nuclear localization signals and are predominantly present in the nucleus during steady state, where they function in various RNA metabolic processes including transcription, splicing, nuclear export, and 3’ end processing ^22,23^, and more recently have been implicated in DNA replication ^24^. A direct connection between chromosome territory associated C0T-1 RNA and hnRNPs was established by the observation that siRNA mediated depletion, or forced expression of dominant interfering mutants, of HNRNPU results in dissociation of C0T-1 RNA from chromosome territories ^13,20,21^. In addition, recent studies have proposed that abundant nuclear proteins such as HNRNPU nonspecifically interact with “RNA debris” that creates a dynamic nuclear mesh that regulates interphase chromatin structure ^12,18^. Here, we used publicly available <u>e</u>nhanced <u>C</u>ross <u>L</u>ink Immuno-<u>P</u>recipitation (eCLIP) data from ENCODE ^25^ to identify RBPs, including HNRNPA1, HNRNPC, HNRNPL, HNRNPM, HNRNPU, and HNRNPUL1, that interact with the known ASAR lncRNAs. We also found that an ∼7 kb region within the ∼185 kb ASAR6-141 lncRNA has a surprisingly high density of interacting RBPs, and that genetic deletion of this ∼7kb “RBP domain” from the endogenous ASAR6-141 locus results in a delayed replication timing phenotype that is comparable to deletion of the entire ∼185 kb ASAR6-141 transcribed region located on human chromosome 6. In addition, ectopic integration of transgenes, expressing the ∼7 kb RBP domain, into autosomes or into the inactive X chromosome results in retention of the RNA within the chromosome territory of the integrated chromosomes, and results in DRT/DMC, and chromosome structure instability in *cis*. Taken together, these results indicate that the ∼7 kb RBP domain represents a critical region within ASAR6-141 that controls replication timing of human chromosome 6.

Because ASARs were identified as *cis-*acting elements that control chromosome-wide replication timing, we tested if ASAR associated RBPs also control replication timing. We found that shRNA mediated depletion of 9 different ASAR-associated RBPs (HNRNPA1, HNRNPC, HNRNPL, HNRNPM, HNRNPU, HNRNPUL1, HLTF, KHSRP or UCHL5) dramatically altered the normally synchronous replication timing program of all autosome pairs, recapitulating the effect of individual ASAR knockouts on a genome-wide scale. These results demonstrate the role that ASARs play during the temporal order of genome-wide replication, and suggests that ASAR lncRNAs serve as essential RNA scaffolds for the assembly of multiple hnRNP/RBP complexes that function to maintain the structural integrity of each mammalian chromosome.

## Results

All of the eight genetically validated ASARs are subject to AEI and express lncRNAs that remain associated with the chromosome territories where they are transcribed ^4–7^. Examples of mono-allelic expression and chromosome territory localization of ASAR6-141 lncRNA are shown in Figure 1A and 1B, where ASAR6-141 lncRNA is localized within one of the chromosome 6 territories in two different cell types, male HTD114 cells and female GM12878 cells, respectively. We note that the size of the RNA hybridization signals detected by all ASAR probes are variable, ranging in size from large clouds that occupy entire chromosome territories, with similarity in appearance to XIST RNA “clouds” (Fig. 1B), to relatively small sites of hybridization that remain tightly associated with the expressed alleles ^4–7^.

**Figure 1.**
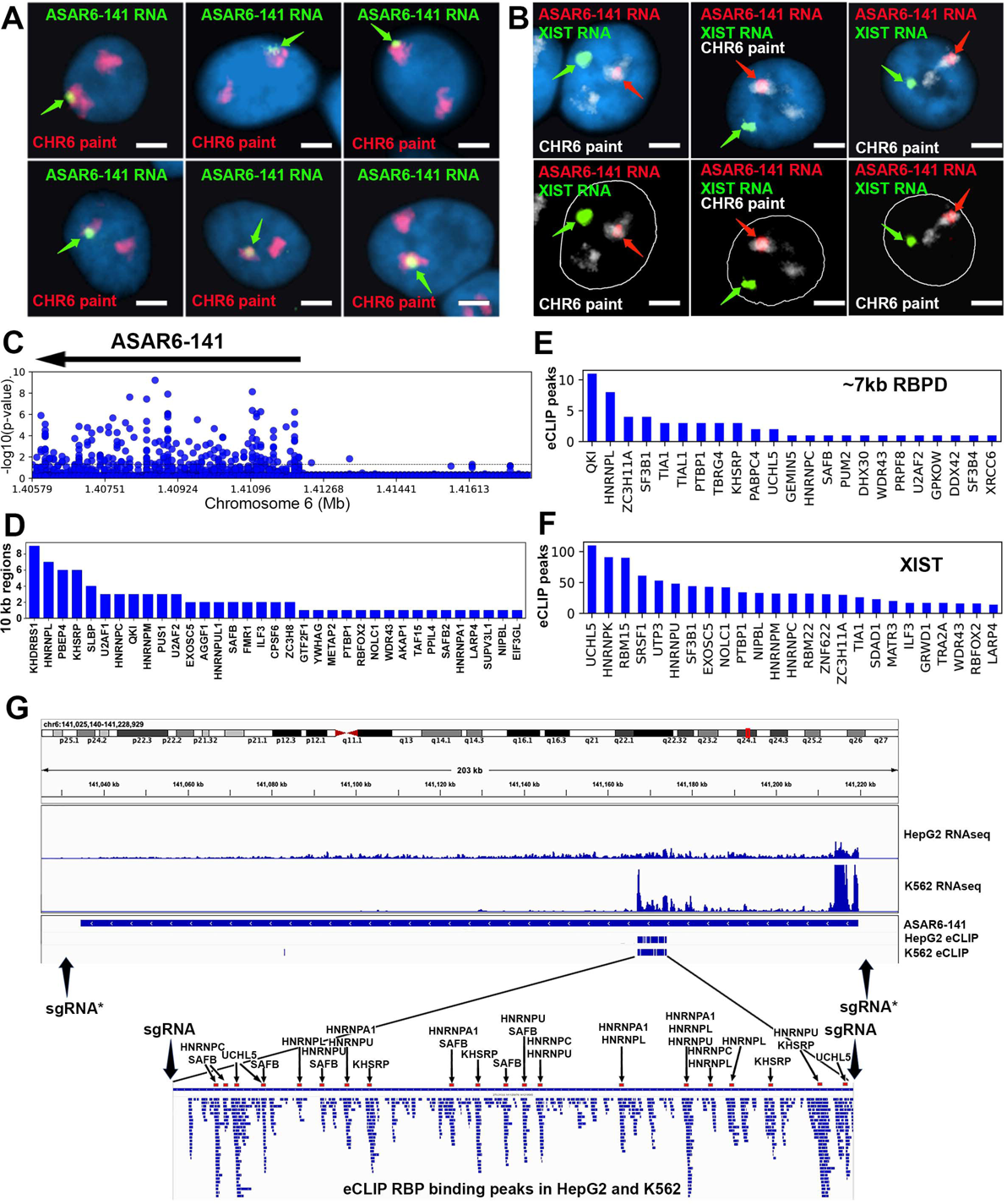
RNA binding proteins interact with an ∼7 kb domain within ASAR6-141 RNA. A) RNA-DNA FISH images of ASAR6-141 expression in six individual HTD114 cells. ASAR6-141 RNA (green; green arrows) and chromosome 6 DNA (chromosome paint, red), and DNA was stained with DAPI. B) RNA-DNA FISH of ASAR6-141 RNA (red, red arrows), CHR6 DNA (chromosome paint, white), and XIST RNA (green, green arrows) visualized within individual GM12878 cells, top and bottom panels represent the same three cells with the nuclear outline drawn in white. C) RNA-protein interaction data (eCLIP from ENCODE) for 120 RBPs expressed in K562 cells for the genomic region that contains ASAR6-141, and ∼200 kb of upstream non-transcribed DNA (chromosome 6 140.8 mb-141.6 mb). Each point represents the FDR-corrected p-value of the z-score of log2-ratio between eCLiP vs control for each RBP within a 10kb sliding window. D) Number of 10kb regions with significant enrichments of eCLiP reads vs control within ASAR6-141. E) Histogram of eCLiP peaks count per RBP within ASAR6-141 7kb RBPD. F) Histogram of eCLiP peaks count per RBP within lncRNA XIST. G) Genome browser view of ASAR6-141 with RNA-seq expression and eCLiP peaks shown in K562 and HepG2. The zoomed in view shows the ∼7kb RBPD with the location of the peaks of eCLIP reads that map within the region (see Figure 1-source data 1). The eCLIP peaks for the RBPs used in the shRNA knockdown experiments are indicated and highlighted with arrows and red bars. The location of sgRNAs are shown with arrows (see Figure 1-source data 3), and the asterisks mark the sgRNAs from ^5^.

We first sought to identify RBPs that interact with the known ASAR lncRNAs. Using publicly available eCLIP data for 150 RBPs in K562 and HepG2 cells (ENCODE; ^25,31^), we identified RBP interactions within 4 ASARs (ASAR1-187, ASAR6-141, ASAR8-2.7, and ASAR9-23). Utilizing the eCLIP peaks previously identified by ENCODE ^25^ we found significant RBP binding peaks for 99 different RBPs that map within the four ASAR lncRNAs (Figure 1-source data 1). We also utilized a “region-based” method ^26^ to identify significant enrichments of eCLiP reads against matched controls in 10 kb windows across the four ASAR loci (Figure 1-source data 2). An example of this analysis is shown in Figure 1C, where ASAR6-141 displays significant RBP interactions for 35 different RBPs across the transcribed region. Given the contiguous expression (RNAseq from total RNA) across the ∼185 kb ASAR6-141 locus in both K562 and HepG2 cell lines (Fig. 1G) and the abundant eCLIP reads from 35 different RBPs in 10 kb windows across the ASAR6-141 lncRNA (Fig. 1C and 1D; see Figure 1-source data 2), we were surprised to detect an ∼7 kb region that contained virtually all of the significant eCLIP peaks in both cell lines (Fig.1G; and Figure 1-source data 1). The zoomed in view in Figure 1G shows alignment of 58 independent eCLIP peaks of RBP interactions within the ∼7kb <u>R</u>NA <u>B</u>inding <u>P</u>rotein <u>D</u>omain (RBPD). Figure 1E shows the 23 RBPs with the most abundant peaks of eCLIP reads within the ∼7kb RBPD. The eCLIP peaks associated with the RBPs chosen for shRNA knockdown experiments (see below) are highlighted in the zoomed in view in Figure 1G.

### Disruption of the ∼7kb RBPD within ASAR6-141 results in delayed replication

To determine if the ∼7kb RBPD within the ASAR6-141 lncRNA contains replication timing activity, we used CRISPR/Cas9 mediated engineering to delete the ∼7 kb sequence from the endogenous locus on human chromosome 6. For this analysis we designed single guide RNAs (sgRNAs) to unique sequences on either side of the ∼7kb RBPD (see Fig. 1G; and Figure 1-figure supplement 1). We expressed these sgRNAs in combination with Cas9 in human HTD114 cells and screened clones for deletion of the ∼7kb RBPD (see Figure 1-figure supplement 1). We chose HTD114 cells for this analysis because they maintain mono-allelic expression and asynchronous replication of both imprinted and random monoallelic genes, they have a stable karyotype that does not change significantly following transfection, drug selection and sub-cloning, and knockouts of the eight known ASARs, including ASAR6-141, result in DRT/DMC and chromosome structure instability in *cis* ^4–9,27,28^. Because ASAR6-141 expression is mono-allelic in the HTD114 cells (see Fig. 1A; and ^5^), we isolated clones that had heterozygous or homozygous deletions of the ∼7kb RBPD. We determined which alleles were deleted based on retention of different base pairs of a heterozygous SNP located within the ∼7kb RBPD (see Figure 1-figure supplement 1 and Figure 1-source data 3).

To assay replication timing of homologous chromosome pairs we quantified DNA synthesis in mitotic cells using a BrdU terminal label assay (Fig. 2A; and ^29,30^). In addition, the HTD114 cells contain a centromeric polymorphism on chromosome 6, which allows for an unambiguous distinction between the two chromosome 6 homologs (^7–9^; also see Fig. 2B). For simplicity, we refer to the chromosome 6 with the larger centromere as CHR6A and to the chromosome 6 with the smaller centromere as CHR6B. From our previous studies we knew that the chromosome 6 with the smaller centromere is linked to the expressed allele of ASAR6-141 ^5^. Figure 2B and 2C shows the replication timing analysis on a mitotic cell where the ∼7kb RBPD was deleted from the expressed allele (CHR6B). Note that CHR6B contains more BrdU incorporation than CHR6A, indicating later replication of CHR6B (Fig. 2D). Quantification of the BrdU incorporation in CHR6B and CHR6A in multiple cells indicated that deletion of the ∼7kb RBPD from the expressed allele resulted in a significant delay in replication timing (Fig. 2E). This is in contrast to cells prior to deletion, or in cells containing a deletion of the ∼7kb RBPD from the silent allele (CHR6A), where the BrdU incorporation is comparable between CHR6A and CHR6B (Fig. 2E). In addition, an asynchronous replication pattern was also present in cells with homozygous deletions of the 7kb RBPD, with CHR6B representing the later replicating allele (Fig. 2E). Furthermore, we found that the delayed replication associated with deletion of the ∼7kb RBPD on CHR6B is comparable to the delayed replication associated with deletion of the entire ASAR6-141 locus from the same chromosome (i.e. CHR6B; Fig. 2E; also see ^5^). These results indicate that the ∼7kb RBPD contains functional sequences for controlling replication timing, and further supports the conclusion that the expressed allele but not the silent allele controls replication timing of human chromosome 6 ^5^.

**Figure 2.**
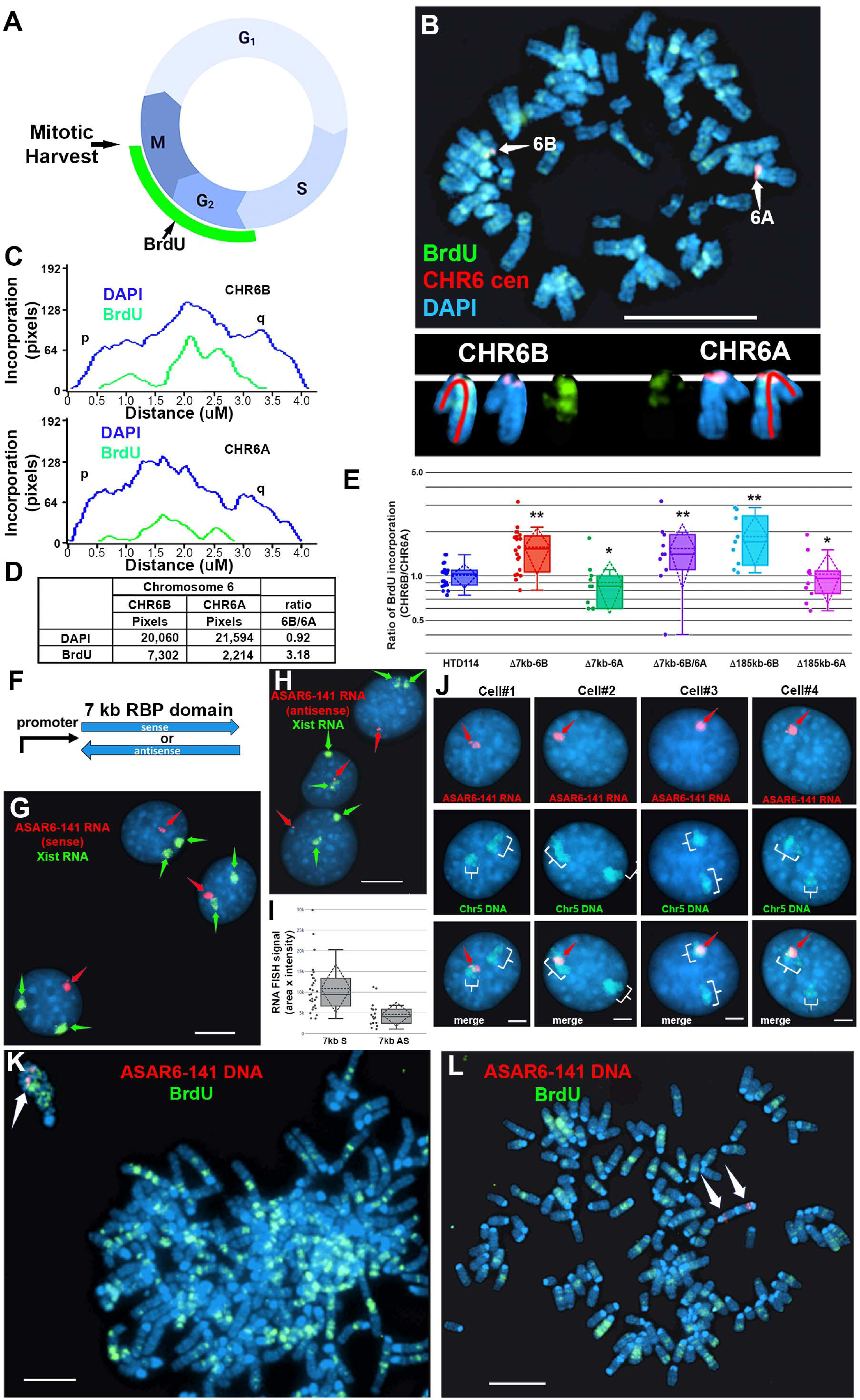
Replication timing in cells with ASAR6-141 *∼7kb* RBPD deletion or ectopic integration. A) Schematic illustration of the BrdU terminal label protocol ^30^. Cells were treated with BrdU (green) for 5 hours and then harvested for mitotic cells. B) BrdU incorporation in HTD114 cells containing a heterozygous deletion of the ∼7kb RBPD. Cells containing a deletion of the ∼7kb RBPD from the expressed allele of ASAR6-141 were exposed to BrdU, harvested for mitotic cells, and subjected to DNA FISH using a chromosome 6 centromeric probe (red; B). The larger centromere resides on the chromosome 6 with the silent ASAR6-141 allele (CHR6A), and the smaller centromere resides on the chromosome 6 with the expressed ASAR6-141 allele (CHR6B). DNA was stained with DAPI (blue). C and D) DAPI staining and BrdU incorporation were quantified by calculating the number of pixels (area x intensity) and displayed as a ratio of BrdU incorporation in CHR6B divided by the BrdU incorporation in CHR6A. E) Quantification of BrdU incorporation in multiple cells with heterozygous (Δ7kb-6B, expressed allele; or Δ7kb-6A, silent allele) or homozygous (Δ7kb-6B/6A) deletions of the ∼7kb RBPD. Also shown is the quantification of BrdU incorporation in heterozygous deletions of the entire ∼185 kb ASAR6-141 gene from chromosome 6B (Δ185-6B) or 6A (Δ185-6A) ^5^. Box plots indicate mean (solid line), standard deviation (dotted line), 25th, 75th percentile (box) and 5th and 95th percentile (whiskers) and individual cells (single points). P values were calculated using the Kruskal-Wallis test ^43^. Values for individual cells are shown as dots. **P = 0.00027 and *P = 0.03. F) Schematic view of the ∼7kb RBPD in both the sense and antisense orientation. The promoter used to drive expression is from ASAR6 ^9^. G and H) Two color RNA FISH assay for expression of the ∼7kb RBPD transgenes. Individual clones were screened for RNA expressed from the ASAR6-141 (∼7kb RBPD) transgenes (red, arrows), and RNA hybridization using an Xist probe (green, arrows) was used as positive control. G and H represent examples of cells containing the sense and antisense transgenes, respectively. I) Quantitation of the size (pixels = area x intensity) of the RNA FISH signals expressed from the 7 kb sense (7 kb S) or antisense (7 kb AS) transgenes are shown. Box plots indicate mean (solid line), standard deviation (dotted line), 25th, 75th percentile (box) and 5th and 95th percentile (whiskers) and individual cells (single points). P value of 0.006 was calculated using the Kruskal-Wallis test ^43^. Values for individual cells are shown as dots. J) RNA from the ∼7kb RBPD remains localized to mouse chromosomes. RNA-DNA FISH using the ∼7kb RBPD as RNA FISH probe (red, arrows), plus a mouse chromosome 5 paint to detect chromosome 5 DNA (green, brackets). K) Cells containing the sense ∼7kb RBPD transgene integrated into mouse chromosome 5 were exposed to BrdU (green), harvested for mitotic cells, and subjected to DNA FISH using the ∼7kb RBPD (red, arrows). Note that the chromosome that contains the transgene shows delayed mitotic condensation and more BrdU incorporation than any other chromosome within the same cell. L) Cells containing the antisense ∼7kb RBPD transgene integrated into a mouse chromosome were exposed to BrdU (green), harvested for mitotic cells, and subjected to DNA FISH using the ∼7kb RBPD (red). Note that the chromosome that contains the transgene shows normal mitotic condensation and only a small amount of BrdU incorporation. Scale bars are 10 uM (B, G, H) 5 uM (K, L) and 2 uM (J).

### Ectopic integration of transgenes expressing the ∼7kb RBPD into autosomes

To determine if the ∼7 kb RBPD is sufficient to induce delayed replication in an ectopic integration assay, we generated two transgenes containing the ∼7kb RBPD, one in the sense orientation and one in the antisense orientation with respect to the promoter (Fig. 2F). These transgenes were introduced into mouse cells and individual clones isolated. We initially screened individual clones for expression of the transgenes using RNA FISH with the ∼7kb RBPD as probe. Because the mouse cells that we used are female, we included an RNA FISH probe for Xist to serve as positive control for the RNA FISH assays. This two-color RNA FISH assay allowed us to identify clones with single sites of transgene expression, and allowed us to directly compare the RNA FISH signals from the transgenes to that of the endogenous Xist RNA expressed from the inactive X chromosome. Figure 2 shows examples of the RNA FISH signals in cells expressing either the sense (Fig. 2G) or antisense (Fig. 2H) transgenes. Large clouds of RNA FISH hybridization signals that are comparable to Xist RNA hybridization signals were detected in 4 out of 12 clones expressing the sense transgene (Fig. 2G). In contrast, 0 out of 12 clones containing the antisense transgene expressed RNA FISH hybridization signals comparable to Xist RNA signals, but instead only small pinpoint sites of RNA FISH hybridization were detected (Fig. 2H). Quantitation of the nuclear area occupied by the sense and antisense ∼7 kb RNAs is shown in Figure 2I. Note that the mouse cells are tetraploid for the X chromosome and contain two Xist RNA hybridization signals, representing two inactive X chromosomes.

To identify the mouse chromosomes that contain the transgenes, we used DNA FISH using the ∼7kb RBPD DNA as probe in combination with the inverted DAPI banding pattern to identify the mouse chromosomes in metaphase spreads. We then confirmed the identity of the mouse chromosomes using DNA FISH on metaphase spreads using the ∼7kb RBPD in combination with BAC probes from the chromosomes of interest. For example, clone ∼7kb(+)A5, which contains the sense-strand transgene integrated in mouse chromosome 5 is shown in Figure 2-figure supplement 1A. Next, to determine if the RNA expressed from the ∼7kb RBPD sense strand transgene is retained within the chromosome territory that expresses the transgene, we used RNA-DNA FISH with the ∼7kb RBPD as RNA probe plus a chromosome paint to detect mouse chromosome 5 DNA in clone ∼7kb(+)A5. Figure 2J shows that the ∼7kb RBPD RNA is localized within one of the chromosome 5 DNA hybridization signals, indicating that the ∼7kb RBPD RNA is retained within the chromosome 5 territory.

Next to determine if the ∼7kb RBPD transgene alters replication timing of mouse chromosome 5, we visualized DNA synthesis in mitotic cells using the BrdU terminal label assay (see Fig. 2A; and ^29,30^). Cells were exposed to BrdU for five hours, harvested for mitotic cells, and processed for BrdU incorporation and for DNA FISH using the ∼7kb RBPD transgene to identify the integrated chromosome. Figure 2K shows an example of this replication-timing assay in a mitotic cell containing the ∼7kb RBPD sense strand transgene integrated into mouse chromosome 5. Note that the chromosome containing the ∼7kb RBPD transgene is delayed in mitotic condensation and contains more BrdU incorporation than any other chromosome within the same cell. In contrast, integration of the transgene with the ∼7kb RBPD in the antisense orientation did not result in delayed condensation nor in delayed replication of mouse chromosomes (Fig. 2L).

### Depletion of RBPs disrupts the chromosome territory localization of ASAR lncRNAs

The results described above identified 99 RBPs expressed in two different cell lines that interact with four different ASAR lncRNAs (see Figure 1-source data 1). Next, to determine if the ASAR associated RBPs function in the chromosome territory localization of ASAR lncRNAs, we analyzed ASAR lncRNA localization in cells with shRNA depletion of 14 different RBPs. Figure 3A-J shows examples of antibody staining for HNRNPU, HNRNPUL1, HNRNPC, HNRNPM, and HLTF in K562 cells expressing either empty vector or vectors expressing shRNAs to individual RBPs. Quantitation of the immunofluorescence for each RBP in empty vector versus shRNA expressing cells, indicated a significant depletion of each RBP (Fig. 3K; also see Figure 3-figure supplement 1 for Western blots).

**Figure 3.**
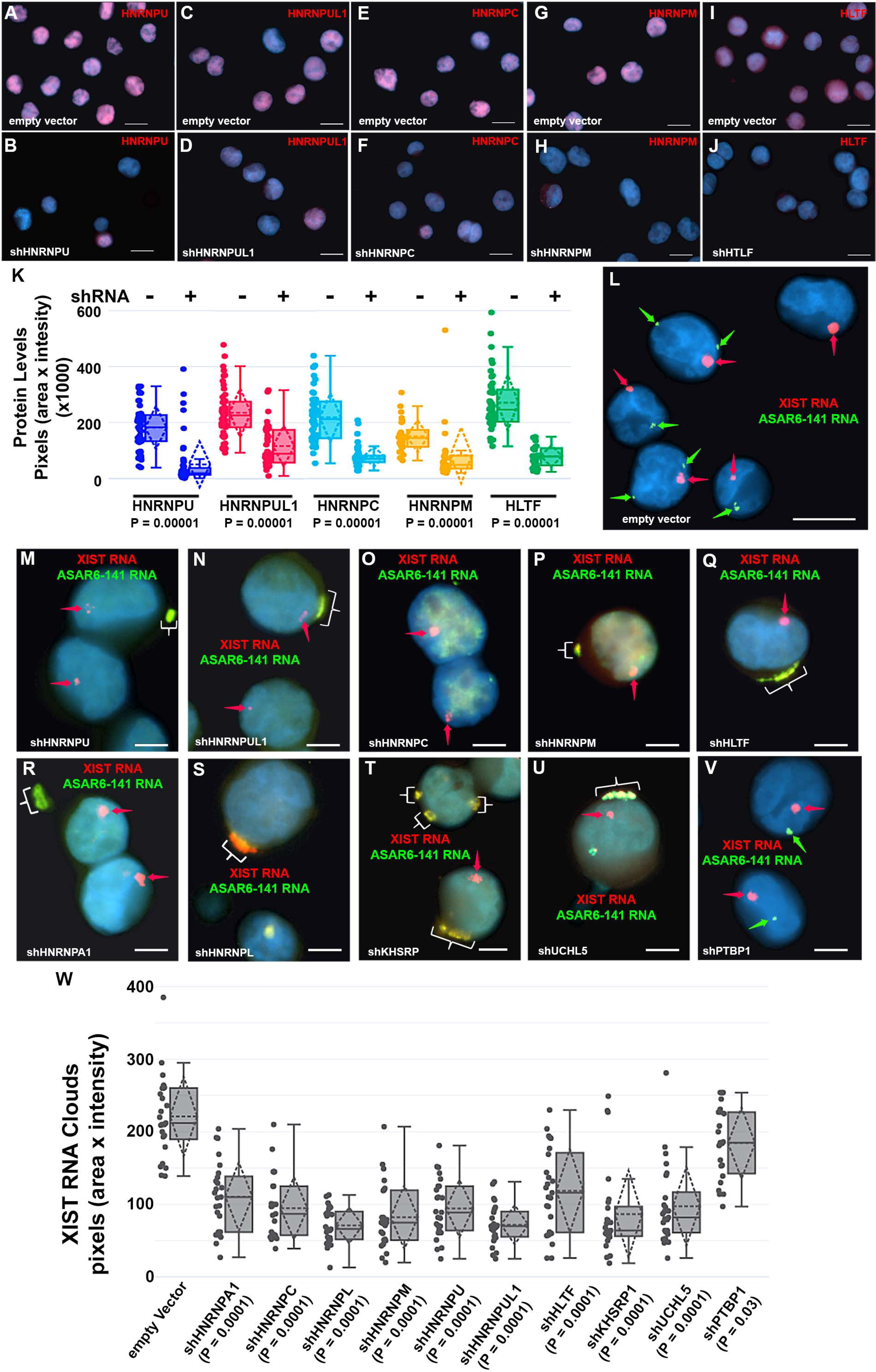
Depletion of RBPs results in disruption of the chromosome territory localization of ASAR6-141 RNA. A-J) shRNA mediated depletion of RBPs. K562 cells were transfected with empty vector (A, C, E, G, and I) or vectors expressing shRNAs directed against HNRNPU (B), HNRNPUL1 (D), HNRNPC (F), HNRNPM (H), or HLTF (J). K) Cells were stained with the appropriate antibodies and quantitation of each RBP was determined in >25 individual cells. Box plots indicate mean (solid line), standard deviation (dotted line), 25th, 75th percentile (box), 5th and 95th percentile (whiskers), and individual cells (single points). P values were calculated using the Kruskal-Wallis test ^43^. L-V) RNA-DNA FISH for ASAR6-141 (green) and XIST (red) RNA. K562 cells were transfected with empty vector (L) or vectors expressing shRNAs against HNRNPU (M), HNRNPUL1 (N), HNRNPC (O), HNRNPM (P), HLTF (Q), HNRNPA1 (R), HNRNPL (S), KHSRP (T), UCHL5 (U), or PTBP1 (V). The red arrows mark the RNA FISH signals for XIST. The brackets mark cytoplasmic regions that hybridized to both RNA FISH probes. DNA was stained with DAPI, and Bars are 10 *u*M (A-J and L) or 5 *u*M (M-V). W) Quantitation of the XIST RNA FISH signals. K562 cells transfected and processed for RNA FISH as in L-V were analyzed for quantitation of XIST RNA cloud size (pixels: area X intensity) in >25 individual cells. Box plots indicate mean (solid line), standard deviation (dotted line), 25th, 75th percentile (box), 5th and 95th percentile (whiskers) and individual cells (single points). P values were calculated using the Kruskal-Wallis test ^43^.

Next, to determine if the nuclear localization of ASAR6-141 lncRNA was affected by RBP depletions we used RNA FISH assays in K562 cells expressing shRNAs against the 14 RBPs. For this analysis we used RNA FISH to detected ASAR6-141 lncRNA, and because K562 cells are female we also included a probe for XIST RNA as positive control for the RNA FISH. Figure 3L shows the expected RNA FISH patterns for both ASAR6-141 and XIST lncRNAs in cells expressing empty vector. In contrast, cells expressing shRNAs directed at HNRNPU, HNRNPUL1, HNRNPC, HNRNPM, HLTF, HNRNPA1, HNRNPL, KHSRP, or UCHL5 showed a nearly complete absence of the highly localized nuclear RNA hybridization signals for ASAR6-141 lncRNA, and the appearance of large punctate cytoplasmic hybridization signals (Fig. 3M-U). We also noticed an overall increase in a diffuse ASAR6-141 lncRNA nuclear and cytoplasmic hybridization signals. In contrast, cells expressing shRNAs directed at PTBP1, PTBP2, or MATR3 showed only the typical nuclear localized hybridization signals for both ASAR6-141 and XIST lncRNAs (Fig. 3V; and Figure 3-figure supplement 2A-D). In addition, localized nuclear XIST RNA hybridization signals were detected in all shRNA treated cells (Fig. 3M-V and Figure 3-figure supplement 2A-D). However, we detected cytoplasmic foci of XIST RNA hybridization that colocalized with the ASAR6-141 lncRNA, and a significant decrease in the size of the nuclear XIST RNA hybridization signals (Fig. 3W). These observations indicate that at least some of the XIST RNA was dissociated from the inactive X chromosome and colocalized with ASAR6-141 lncRNA in large cytoplasmic foci.

### Ectopic integration of the ∼7kb RBPD transgene into the inactive X chromosome

The results described above indicate that shRNA depletion of 9 different RBPs resulted in dissociation of ASAR6-141 lncRNA from its parent chromosome, and resulted in a decreased nuclear XIST RNA hybridization signal with concomitant colocalization of both RNAs in cytoplasmic foci (see Fig. 3), suggesting that ASAR6-141 and XIST lncRNAs are retained on their respective chromosome territories via similar mechanisms, which is supported by the large overlap in RBPs associated with both RNAs (Fig. 1E and 1F; and Figure 1-source data 1). In addition, ectopic integration of *Xist* transgenes into autosomes has been an instrumental tool in characterization of *Xist* functions, including the ability to delay replication timing and induce gene silencing ^31–33^. Therefore, we next sought to determine the consequences of ectopic integration of the ASAR6-141 ∼7kb RBPD transgene into the inactive X chromosome. The ectopic integration assays described above involved random integration of transgenes followed by an RNA FISH screen for clones expressing the ∼7kb RBPD, which included an RNA FISH probe for Xist RNA as positive control. Therefore, to identify transgene integrations into the inactive X chromosome, we simply screened an additional 50 clones for colocalization of RNA FISH hybridization signals for both Xist and the ∼7kb RBPD transgene. Figure 4A shows this RNA FISH assay on cells where the Xist and ∼7kb RBPD hybridization signals are colocalized. We note that the intensity of the Xist RNA FISH signal was enhanced in the region where the ∼7kb RBPD hybridization signal was detected (Fig. 4B), suggesting that the presence of the ∼7kb RBPD RNA promoted preferential localization of Xist RNA in the same region of the X chromosome territory that contains the ∼7kb RBPD RNA.

**Figure 4.**
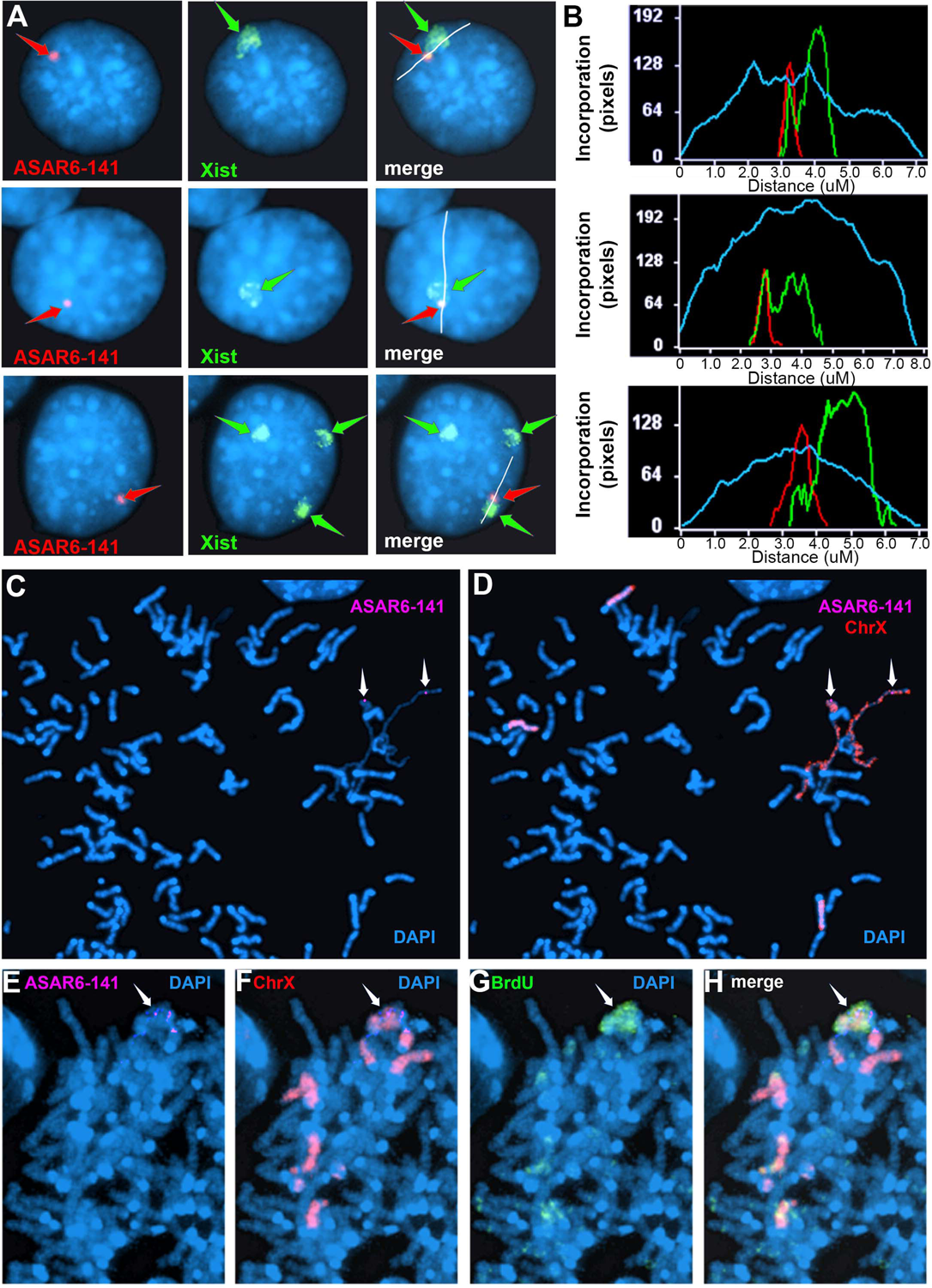
Ectopic integration of the ASAR6-141 *∼7kb RBPD* transgene into an inactive X chromosome. A) Two color RNA FISH assay for expression of the sense ∼7kb RBPD transgene plus Xist RNA. Examples of three different cells showing colocalization of the ASAR RNA and Xist RNA. Individual cells were processed for expression of the sense strand ∼7kb RBPD transgene (left panels, red, arrow) in combination with RNA FISH for Xist (middle panels, green, arrow). The right panels show the merged images. The white lines show the path for the pixel intensity profiles used for panel B. B) Pixel intensity profiles across the Xist RNA hybridization domain showing enhanced signal intensity over the ∼7kb RBPD RNA hybridization domain. C and D) Delayed mitotic chromosome condensation of the X chromosome that contains the ∼7kb RBPD transgene. An example of a mitotic spread processed for DNA FISH using the ∼7kb RBPD as probe (magenta, panel C) plus a mouse X chromosome paint (red, panel D). The arrows mark the location of the transgene hybridization signal. E-H) Cells containing the ∼7kb RBPD transgene integrated into the mouse X chromosome were exposed to BrdU (G, green), harvested for mitotic cells, and subjected to DNA FISH using the ∼7kb RBPD (E, magenta) plus a mouse X chromosome paint (F, red). The merged images are shown in panel H.

To confirm that the mouse X chromosome contains the transgene, we used DNA FISH using the ∼7kb RBPD DNA as probe in combination with an X chromosome paint probe on metaphase spreads. An example of this analysis is shown in Figure 2-figure supplement 1B-C, and indicates that the ∼7kb RBPD transgene is integrated into an X chromosome. During this analysis we detected DMC on X chromosomes, and the X chromosomes with DMC contain the ∼7kb RBPD transgene (Figs. 4C and 4D).

Next, to determine if the ∼7kb RBPD transgene alters replication timing of the inactive X chromosome, we visualized DNA synthesis in mitotic cells using our BrdU terminal label assay (see Fig. 2A; and ^29,30^). Cells were exposed to BrdU for five hours, harvested for mitotic cells, and processed for BrdU incorporation and for FISH using the ∼7kb RBPD transgene plus an X chromosome paint probe to identify the integrated chromosome. Figure 4E-4H shows an example of delayed replication and delayed mitotic condensation of the X chromosome in a mitotic cell containing the ∼7kb RBPD sense transgene integrated into the inactive X chromosome. We conclude that integration of the 7kb RBPD transgene is sufficient to cause DRT/DMC on the inactive X chromosome.

### Disruption of the chromosome associated localization of ASAR6 and ASAR1-187 lncRNAs

To determine if the chromosomal localization of additional ASAR RNAs was also affected by RBP knockdowns we carried out RNA-DNA FISH for ASAR1-187 and ASAR6 lncRNAs. For this analysis we used Fosmid probes to detect the ASAR RNAs plus BAC probes to detect the DNA near the ASAR loci located on chromosomes 1 and 6 (see Figure 1-source data 3). Figure 5A-5C shows this RNA FISH assay on K562 cells expressing empty vector, and indicates that the RNA FISH signals for both ASARs are closely associated with the DNA FISH signals on their respective chromosomes, i.e. ASAR1-187 RNA shows close association with the chromosome 1 BAC DNA, and ASAR6 RNA shows close association with the chromosome 6 BAC DNA. In contrast, cells with shRNA depletion of HNRNPA1, HNRNPC, HNRNPL, HNRNPM, HNRNPU, HLTF, KHSRP, or UCHL5 showed a dispersed punctate nuclear pattern for both ASAR RNAs (Fig. 5D-5K). In addition, large punctate cytoplasmic foci were detected, with the two ASAR RNA hybridization signals colocalized.

**Figure 5.**
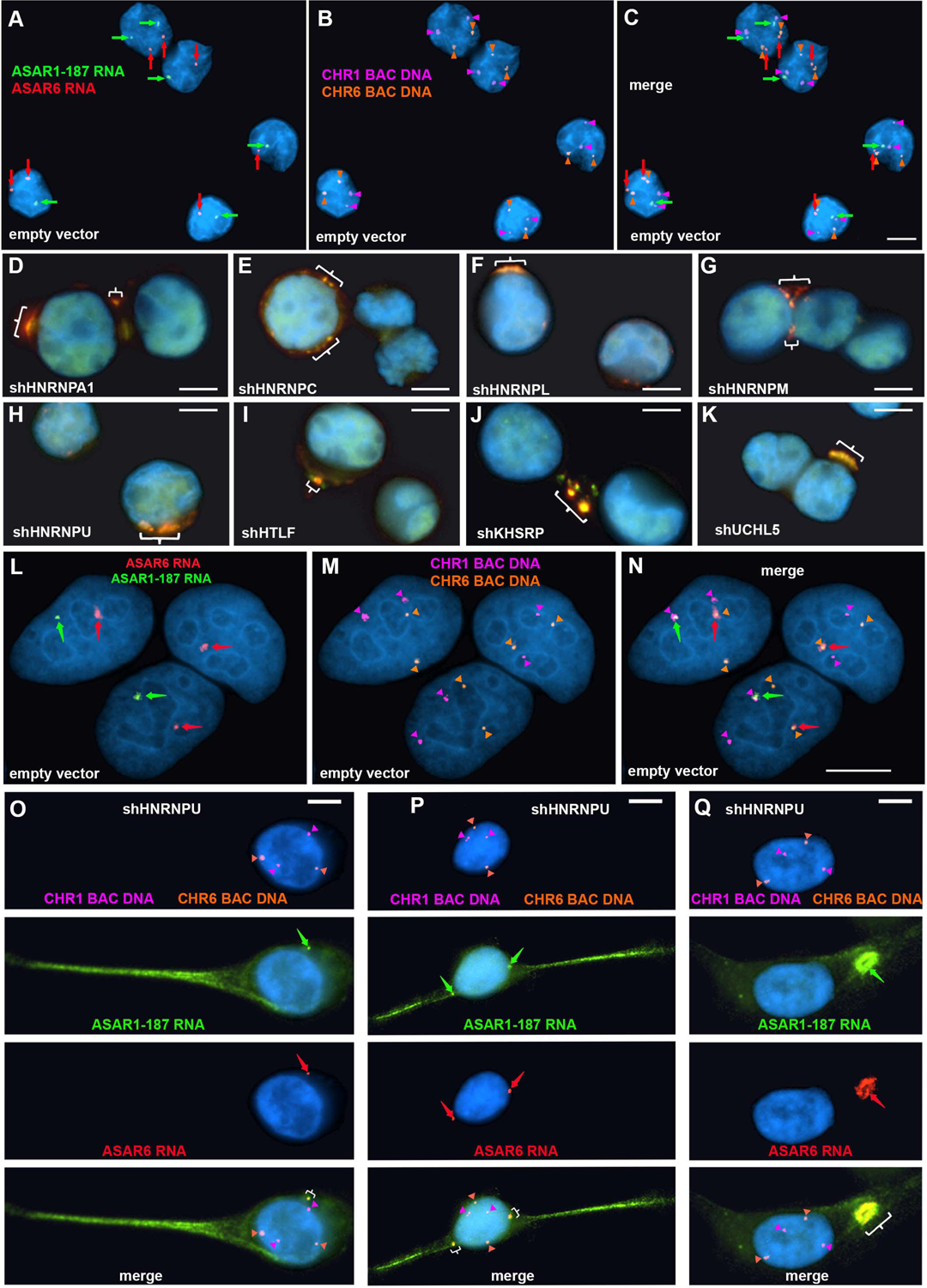
Depletion of RBPs results in disruption of the chromosome territory localization of ASAR RNAs. shRNA mediated depletion of RBPs in K562 cells (A-K) or HTD114 cells (L-Q). Cells were transfected with empty vector (A-C and L-N) or vectors expressing shRNAs directed against HNRNPA1 (D), HNRNPC (E), HNRNPL (F), HNRNPM (G), HNRNPU (H, and O-Q), HLTF (I), KHSRP (J), or UCHL5 (K). Cells were processed for RNA FISH with probes for ASAR6 (red, arrows) and ASAR1-187 (green, arrows). Cells were also processed for DNA FISH using BAC DNA to detect chromosome 1 (CHR1 BAC DNA, magenta) and chromosome 6 (CHR6 BAC DNA, orange; A-C and L-Q). Arrows mark the sites of RNA FISH hybridization and arrowheads mark the sites of DNA hybridization. RNA-DNA FISH for ASAR1-187 (RNA in green, arrows; DNA in magenta and arrowheads), ASAR6 (RNA in red, arrows; DNA in orange and arrowheads). The brackets mark the cytoplasmic regions that hybridized to both RNA FISH probes. DNA was stained with DAPI, and scale bars are 10 *u*M.

Next, to determine if depletion of RBPs would result in loss of the ASAR RNA chromosome localization in a second cell type, we carried out RNA-DNA FISH assays on shRNA depleted HTD114 cells. We chose HTD114 cells because we previously used these cells to detect the chromosome territory localization of all eight ASAR RNAs and to measure the synchronous replication timing of five autosome pairs both before and after ASAR disruptions (^4–8,28^; and see Figure 2). Figure 5L-5N shows the RNA-DNA FISH assay in HTD114 cells expressing empty vector. We again note the tight association of the ASAR RNAs with the genomic loci where they are transcribed. In contrast, cells expressing shRNA against HNRNPU show loss of the chromosome associated RNA hybridization signals for both ASAR RNAs, and instead show diffuse nuclear and cytoplasmic hybridization, with the presence of large punctate cytoplasmic foci that hybridize to both ASAR RNA FISH probes (Fig. 5O-5Q). Similar results were obtained following shRNA depletion of HNRNPA1, HNRNPC, HNRNPL, HNRNPM, HNRNPUL1, KHSRP, HLTF, or UCHL5 (Figure 5-figure supplement 1). In contrast, expression of shRNAs against SAFB, SAFB2, SAFB plus SAFB2, CTCF, or MATR3 did not alter the ASAR1-187 or ASAR6 RNA nuclear hybridization signals in HTD114 cells (Figure 3-figure supplement 2E-J).

### Depletion of RBPs causes asynchronous replication on all autosome pairs

Next, to determine if ASAR associated RBPs function during the normally synchronous replication timing program that occurs on pairs of homologous chromosomes, we analyzed DNA synthesis in HTD114 cells depleted for the 9 RBPs implicated in ASAR RNA localization (HNRNPU, HNRNPA1, HNRNPC, HNRNPL, HNRNPM, HNRNPUL1, KHSRP, HLTF, or UCHL5) and for five RBPs that showed no effect on ASAR RNA localization (MATR3, PTBP1, PTBP2, SAFB, or SAFB2). Using the BrdU terminal label assay ^29,30^ in cells expressing the empty vector, we found that each autosome pair displays a stereotypical and synchronous BrdU incorporation pattern (Fig. 6A). In contrast, following HNRNPU depletion, individual mitotic spreads displayed dramatic differential incorporation of BrdU into pairs of homologous chromosomes, examples of asynchronous replication timing between every pair of autosomes are shown in Figure 6B. Note that each chromosome pair contains dramatic differential BrdU incorporation into chromosome homologs, which represents considerable asynchronous replication timing. Figure 6C and 6D show examples for chromosome pairs 4 and 18 (also see Figure 6-figure supplement 1C). Quantification of the BrdU incorporation in chromosomes 1 and 6 (see Figure 6-figure supplement 1) in multiple cells depleted for HNRNPU, HNRNPUL1, HLTF, KHSRP, UCHL5, or HNRNPA1 indicated that both pairs of chromosomes display significant asynchrony (Fig. 6E and 6F). In addition, the DRT/DMC phenotype was detected in ∼30% of mitotic spreads following shRNA-mediated depletions (two examples are shown in Fig. 6G and 6H; also see Figure 6-figure supplement 2). Furthermore, shRNA-mediated depletion of HNRNPC, HNRNPL and HNRNPM resulted in dramatic alterations in chromosome morphology that included asynchronous replication and abnormal mitotic condensation, including ∼10% of mitotic spreads with one or more chromosomes that displayed the DRT/DMC phenotype (Figure 6-figure supplement 2).

**Figure 6.**
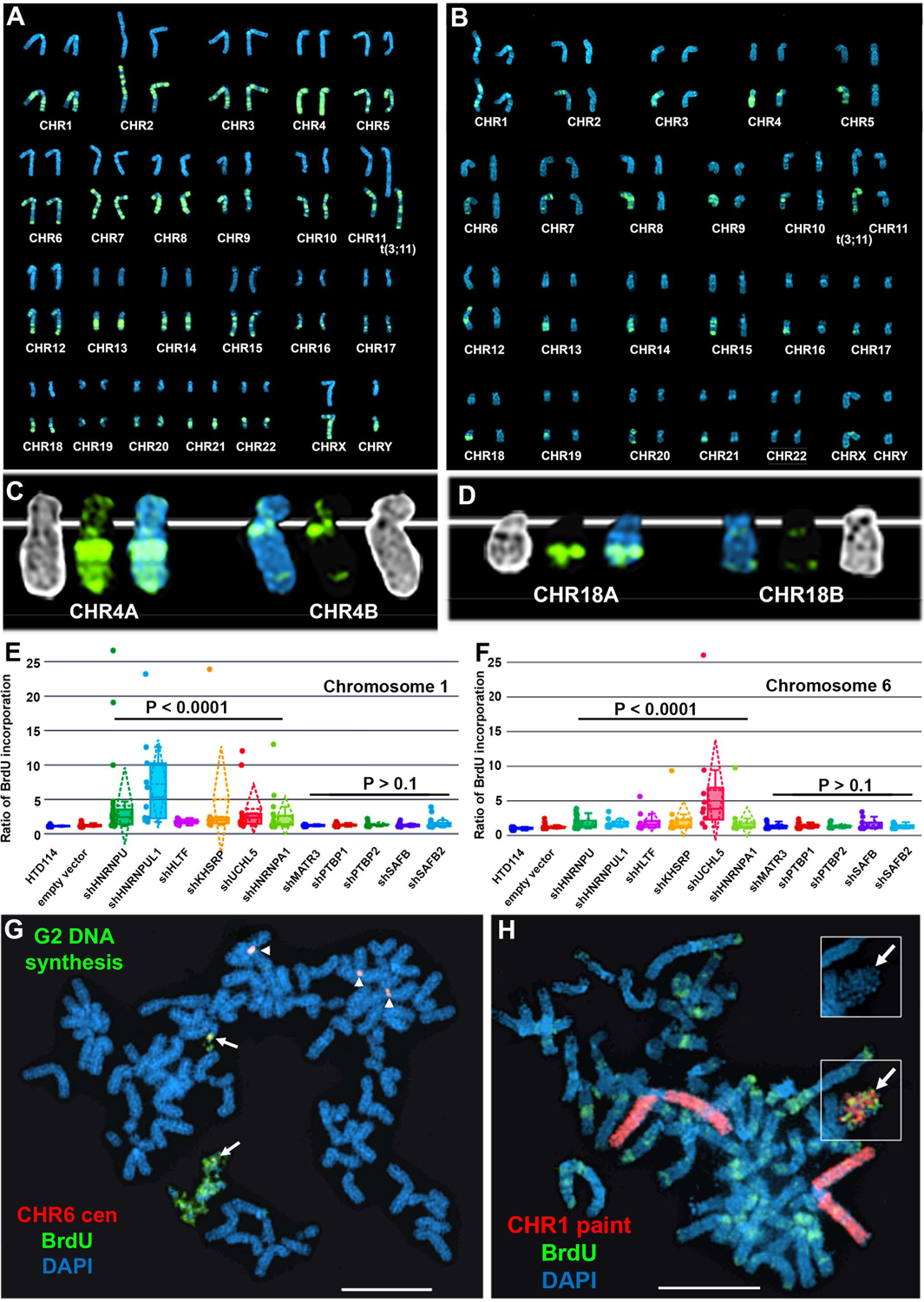
Depletion of RBPs results in asynchronous replication on autosome pairs. A) BrdU incorporation in parental HTD114 cells. Cells were exposed to BrdU for five hours and processed for BrdU incorporation using an antibody against BrdU. This panel represents chromosomes from multiple mitotic spreads showing representative BrdU incorporation in chromosome pairs. Both homologs of autosome pair were captured from the same mitotic cell, and each pair displays a typical BrdU incorporation pattern that is consistent with synchronous replication timing. B) HTD114 cells were transfected with the HNRNPU shRNA expression vector, exposed to BrdU for five hours, and processed for BrdU incorporation. This panel represents chromosomes from multiple mitotic spreads showing representative BrdU incorporation in pairs of autosomes. Both homologs of autosome pairs were captured from the same mitotic cell, and each pair displays a differential BrdU incorporation pattern that is consistent with asynchronous replication timing. C) Shows two chromosome 4 homologs (CHR4A and CHR4B) side by side. D) Shows the BrdU incorporation in chromosome 18 homologs (CHR18A and CHR18B). For each chromosome pair the inverted DAPI staining (black and white), BrdU incorporation (green), and DAPI staining (blue) are shown. E and F) Quantification of BrdU incorporation in multiple cells depleted for HNRNPU, HNRNPUL1, HTLF, KHSRP, UCHL5, HNRNPA1, MATR3, PTBP1, PTBP2, SAFB and SAFB2. Box plots indicate mean (solid line), standard deviation (dotted line), 25th, 75th percentile (box), 5th and 95th percentile (whiskers) and individual cells (single points). P values were calculated using the Kruskal-Wallis test ^43^. G) DRT/DMC on chromosome 6 following HNRNPU depletion. HTD114 cells were transfected with the HNRNPU shRNA expression vector, exposed to BrdU for 5 hours, harvested for mitotic cells, and processed for BrdU incorporation (green) and DNA FISH using a chromosome 6 centromeric probe (red). H) DRT/DMC on chromosome 1 following UCHL5 depletion. HTD114 cells were transfected with the UCHL5 shRNA expression vector, exposed to BrdU for 5 hours, harvested for mitotic cells, and processed for BrdU incorporation (green) and DNA FISH using a chromosome 1 paint as probe (red). The arrow marks the chromosome 1 with DRT/DMC, and the inset shows only the DAPI staining of the chromosome 1 highlighting the DMC.

### C0T-1 DNA hybridizes to ASAR6 RNA

Finally, to determine the relationship between the RNA species detected by C0T-1 DNA and ASAR RNAs in RNA FISH assays, we used human C0T-1 DNA as an RNA FISH probe on mouse cells expressing an ASAR6 BAC transgene ^8^. Because human C0T-1 DNA does not cross hybridize with mouse RNA or DNA in FISH assays ^20^, we were able to detect the chromosome territory localized expression of human ASAR6 RNA expressed from the transgene that is integrated into one of the mouse chromosome 3 homologs using human C0T-1 DNA as RNA FISH probe (Fig. 7A and 7B), indicating that C0T-1 DNA detects ASAR6 RNA.

**Figure 7.**
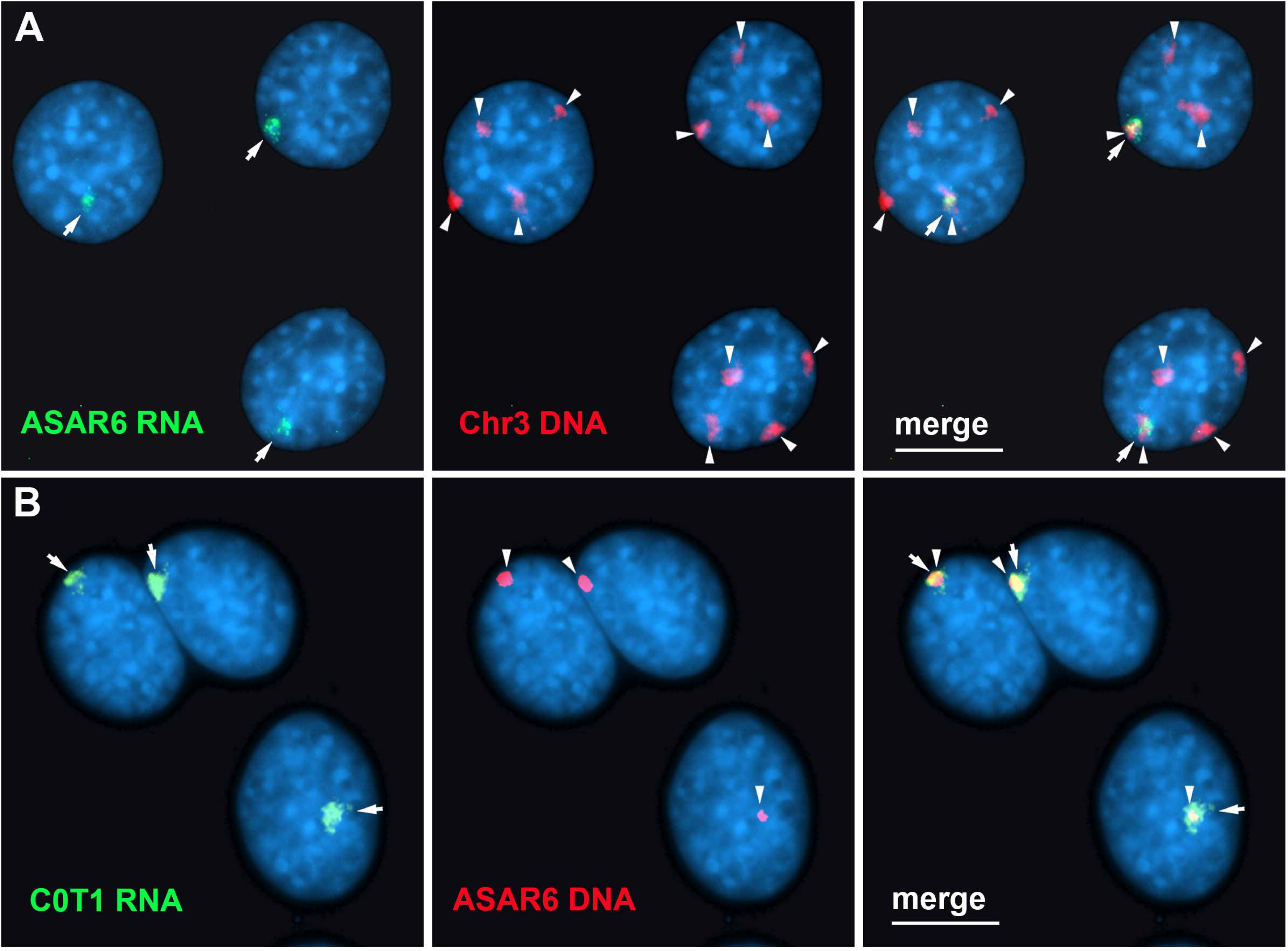
Human C0T-1 DNA detects ASAR6 RNA. A and B) RNA-DNA FISH on mouse cells containing an ASAR6 BAC transgene integrated into mouse chromosome 3 ^8^. A) RNA-DNA FISH using an ASAR6 fosmid probe to detect ASAR6 RNA (green; arrows) plus a mouse chromosome 3 paint probe to detect DNA (red; arrowheads). B) RNA-DNA FISH using human C0T-1 DNA to detect RNA (green; arrows) plus the ASAR6 BAC (red arrowheads) to detect DNA.

## Discussion

We recently identified 68 ASAR candidates, and genetically validated five out of five as controlling replication timing of individual human autosomes, bringing the total of genetically validated ASARs to eight ^4–7^. These eight ASARs (ASAR1-187, ASAR6, ASAR6-141, ASAR8-2.7, ASAR9-23, ASAR9-24, ASAR9-30 and ASAR15) display striking epigenetically controlled differential allelic expression and replication timing, and their RNAs remain localized to their parent chromosome territories ^4–7^. In this report, we used publicly available RNA-protein interaction data from ENCODE to identify RBPs that interact with multiple ASAR lncRNAs. One unanticipated observation that came from this analysis was that an ∼7 kb domain within the ∼185 kb ASAR6-141 lncRNA contained virtually all of the significant peaks of RBP interactions in two different cell lines. Using genetic deletion and ectopic integration assays we found that this ∼7kb RBPD contains functional sequences for controlling replication timing of entire chromosomes in *cis*. We also found that shRNA-mediated depletion of 9 different ASAR-associated RBPs results in loss of the chromosome territory association of multiple ASAR RNAs. In addition, we found that depletion of HNRNPA1, HNRNPC, HNRNPL, HNRNPM, HNRNPU, HNRNPUL1, HLTF, KHSRP, or UCHL5 alters the normally synchronous replication timing that occurs between pairs of homologous chromosomes resulting in the DRT/DMC phenotype, recapitulating the effect of individual ASAR knockouts on a genome-wide scale.

Previous studies have suggested that C0T-1 RNA plays a dynamic structural role that promotes the open architecture of active chromosome territories ^13,20^. C0T-1 RNA is predominantly L1 sequences, and these L1 RNAs remain associated with the chromosome territories where they are transcribed ^20^. Additional studies have proposed that repRNA is an important regulator of the dynamic organization of genomic loci into membrane-less subcompartments with distinct nuclear functions (reviewed in ^19^). Other studies have proposed that HNRNPU nonspecifically interacts with “RNA debris” to create a dynamic nuclear mesh that regulates interphase chromatin structure ^12,18^. One important distinction that can be made between ASAR RNAs and these other hnRNAs (i.e. C0T-1 RNA, repRNA, caRNA, and “RNA debris” ^12,13,18–21^) is that ASARs have been defined genetically, and these other RNAs have not. Thus, we have used both loss of function and gain of function assays to define the roles that ASARs play in chromosome dynamics, which includes control of chromosome-wide replication timing and mitotic chromosome condensation ^4–8^. We also found that C0T-1 DNA, when used as an RNA FISH probe, can detect ASAR6 RNA that is expressed and localized to an individual chromosome territory, indicating that at least some of the RNA FISH signal detected by C0T-1 DNA represents ASAR RNA. Given our recent discovery of ASAR RNAs expressed from every autosome, and encoded by ∼3% of the human genome ^6^, we propose that ASAR RNAs represent functional “chromosome associated RNA” species that control chromosome dynamics within mammalian nuclei. Whether or not there are nuclear functions of other chromosome associated hnRNAs that are independent of ASARs will require genetic and/or functional validation.

Here we identified several hnRNPs (HNRNPA1, HNRNPC, HNRNPL, HNRNPM, HNRNPU, and HNRNPUL1) as important interaction partners for the chromosome territory localization of multiple ASAR RNAs. In addition, we found that the SWI/SNF related transcription factor HLTF, which was recently shown to regulate replication fork reversal, prevent alternative mechanisms of stress-resistant DNA replication, and mediates cellular resistance to replication stress ^34^, is required for the chromosome territory localization of multiple ASAR RNAs and for the synchronous replication of every autosome pair. Furthermore, we found that the KH domain (HNRNP<u>K</u> <u>H</u>omology domain) containing protein KHSRP, and the ubiquitin hydrolase UCHL5 also contribute to the chromosome territory localization of ASAR RNAs. We found that depletion of HNRNPA1, HNRNPC, HNRNPL, HNRNPM, HNRNPU, HNRNPUL1, HLTF, KHSRP, or UCHL5 proteins resulted in a dramatic genome-wide disruption of the normally synchronous replication timing program on all autosome pairs. Taken together our results indicate that the association of ASAR RNAs with multiple hnRNPs in combination with other nuclear RBPs is an essential step in the regulation of the synchronous replication timing program on mammalian chromosomes.

While these studies have demonstrated a role for multiple RBPs in genome-wide chromosome replication timing, one limitation of these studies is that we only tested 14 out of the ∼100 RBPs shown by eCLIP to associate with multiple ASAR RNAs. It is likely that additional insights into the network of RBP/ASAR interactions could be gained by extending our replication timing analyses to knockdowns of additional RBPs. Another limitation of these studies is that all of the RBPs implicated in control of replication timing are known to play various roles in other essential nucleic acid metabolic pathways, including transcription, splicing, mRNA export, and RNA degradation ^22,23^. It is possible that knocking down these RBPs has an indirect effect on replication related to these other functions of the RBPs, rather than a direct effect on ASAR function. However, the RBP knockdowns that disrupt genome-wide synchronous replication also disrupt chromosome association of multiple ASAR RNAs. Our previous work demonstrated that the ability of an ectopically integrated ASAR6 transgene to cause delayed replication timing of a mouse chromosome required the ASAR6 RNA ^9^. Hence, we expect that displacement of ASAR RNA from chromosome territories upon RBP knockdown is a primary cause of the genome-wide replication timing asynchrony, regardless of whether the displacement of ASAR RNA from the chromosome is a direct or indirect effect of RBP knockdown. A final limitation of these studies is that the precise mechanism by which ASARs and RBPs interact to control replication timing remains unknown. One intriguing possibility that is consistent with our data is that the ASAR RNAs serve as scaffolds for the assembly of extensive RNA-protein and protein-protein complexes that form a chromosome-tethered liquid-liquid phase that facilitates the recruitment of the requisite chromatin modifiers and DNA replication machinery. In support of this notion, HNRNPA1 and HNRNPC are components of the 40S hnRNP particle, which forms a nuclear biomolecular condensate ^23^. In addition, HNRNPU and HNRNPUL1 interact with each of the 40S hnRNP particle subunits ^35^, suggesting the possibility of extensive protein-protein interactions. ASAR RNAs normally remain associated exclusively with the chromosome territory from which they are transcribed, and ASAR RNAs on different chromosomes remain separate. RBP knockdowns that disrupt synchronous replication timing also release ASAR RNAs from their chromosome territories, allowing colocalization of ASAR RNAs from different chromosomes (i.e. ASAR6 and ASAR1-187, see Fig. 5O-Q), often in nuclear and/or cytoplasmic foci reminiscent of the mixing of liquid-liquid phases. Additional support for this model is the observation that Xist RNA participates in the assembly of a nuclear heteromeric condensate essential for gene silencing on the inactive X chromosome ^36^. It is worth noting that ectopic integration of Xist transgenes has been a useful assay for characterization of Xist functions, including the ability to delay replication timing and induce gene silencing on autosomes (reviewed ^31^). Thus, our observation that the ∼7kb RBPD RNA induces DRT/DMC on the inactive X chromosome, and associates with the Xist RNA cloud (Fig. 4A and 4B), suggests that the ∼7kb RBPD RNA interferes with Xist RNA function within the heteromeric condensate. One intriguing observation, which is largely ignored by the X inactivation field, is that deletion of the Xist gene on either the active or inactive X chromosomes results in delayed replication timing of the X chromosome ^37,38^. Thus, loss of function mutations of either ASARs or Xist result in a similar chromosome-wide delayed replication timing phenotype. These parallels between ASAR and Xist mutation phenotypes suggest a shared mechanism for controlling replication timing in cis.

The formation of liquid-liquid phases via ASAR RNA-RBP interactions is speculative at this point and remains to be tested directly. A full understanding of the mechanisms by which ASARs control chromosome replication timing will require an extensive biochemical and biophysical characterization of the ASAR RNA-protein and protein-protein complexes and how they interact with the DNA replication machinery.

## Materials and Methods

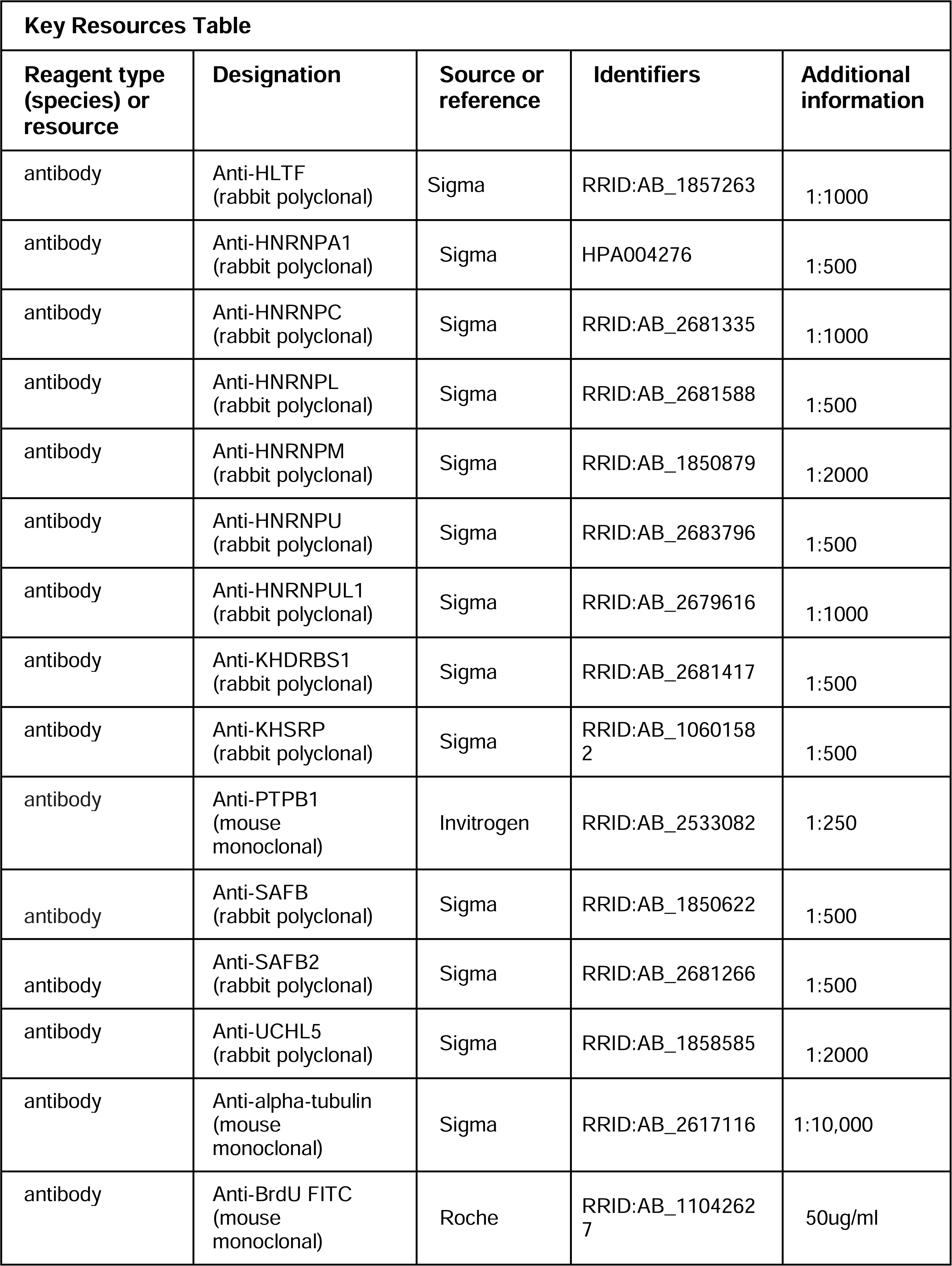
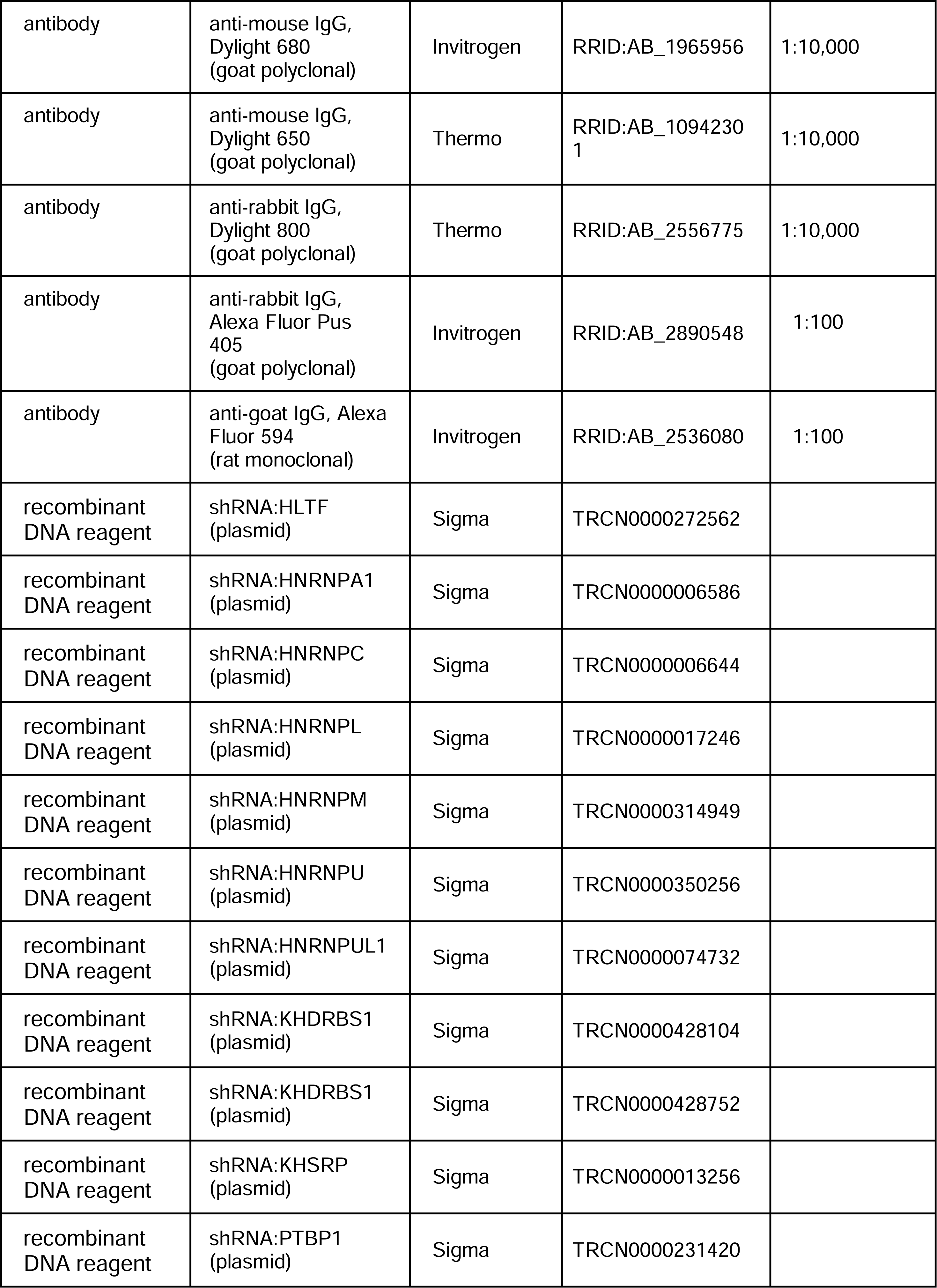
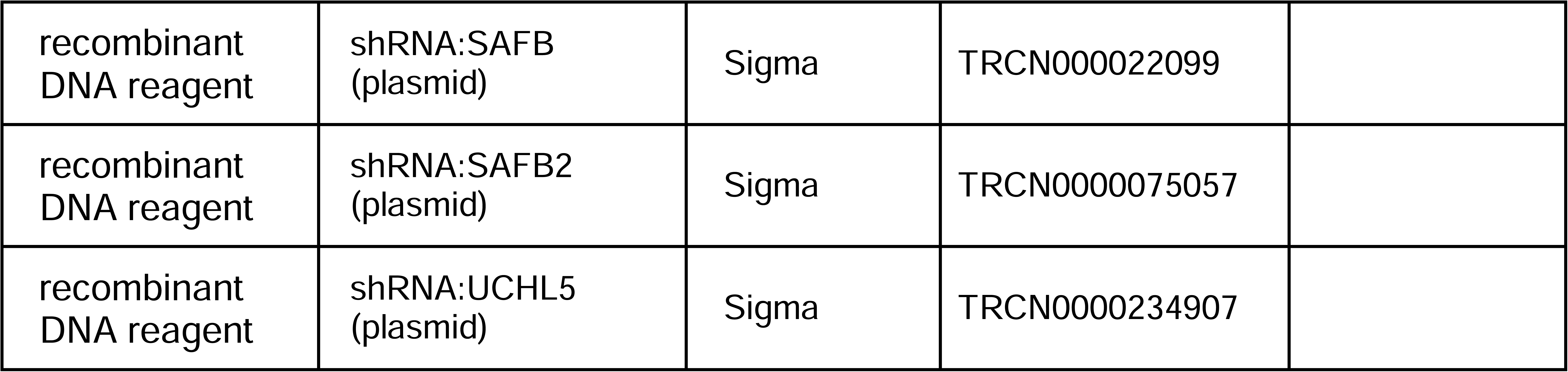

### Cell culture

HTD114 cells are a human *APRT* deficient cell line derived from HT1080 cells ^39^. K562 cells were obtained from ATCC, and cultured in RPMI 1640 (Gibco) supplemented with 10% fetal bovine serum (Hyclone). Mouse C2C12 cells were used for the transgene integration assays, transfected using Lipofectamine 2000 (Invitrogen) according to the manufacturer’s recommendations, and clones containing transgene integrations were isolated following selection in media containing G418. All cells were grown in DMEM (Gibco) supplemented with 10% fetal bovine serum (Hyclone), and were grown in a humidified incubator at 37°C in a 5% carbon dioxide atmosphere.

### eCLIP analysis

eCLIP data was obtained from ^40^. Given that non-coding genes and TLs are expressed at a lower level than coding genes, a sensitive filtering approach was used as recommended by the authors ^40^. We filtered out peaks containing an eCLIP peak p-value greater than p=0.05, and filtered out peaks containing a fold enrichment value less than two.

### Identification of TL

Nuclear-enriched, strand-specific, ribo-minus, total-RNA sequencing libraries were first aligned to the hg19 reference genome using default parameters with STAR ^41^. Next SAMtools ^42^ was used to remove duplicate and low quality (<=MAPQ20) reads. TLs were defined using a strategy of sequential merging of strand-specific, contiguous intergenic reads, as performed in ^6^. Utilizing strand-specific reads, stranded reads separated by 1,000 bp or less were merged to create contigs, and the contigs merged again while allowing for gaps of 7 kb to allow for regions that are not uniquely mappable due to presence of full-length LINE elements. TL were then classified above a minimum cutoff length of 50 kb of strand specific, contiguous expression.

### shRNA depletion of RBPs

Using Lipofectamine 2000 (Invitrogen), according to the manufacturer’s recommendations, we co-transfected HTD114 cells with plasmids encoding puromycin resistance (PGK-PURO) and shRNAs against 14 different RBPs (see Key Resource Table). 24h after transfection, cells were plated in media containing 10 ug/ml puromycin (Millipore/Sigma), and 24h-48h following puromycin selection cells were processed for Western blot, RNA-DNA FISH or for replication timing assays (see below).

### Western blot analysis

Cells were pelleted, rinsed once in PBS, and lysed by resuspension in RIPA Solubilization Buffer (25 mM Tris-HCl pH 7.6, 150 mM NaCl, 5 mM EDTA,1% NP-40 or 1% Triton X-100, 1% sodium deoxycholate, 0.1% SDS). Protein concentration was determined via the Qubit Protein BR Assay kit (Invitrogen, A50669) on a Qubit 4 Fluorometer. Lysates from HTD114 (20 μg) or K562 (25 μg) cell lines in 1X Laemmli Sample Buffer (Bio-Rad, 161-0737) with 5% β-mercaptoethanol were loaded onto 10% Mini-PROTEAN TGX Precast Tris-Glycine-SDS polyacrylamide gels, and run in 1X SDS PAGE running buffer (Bio-Rad, 161-0732) at 100 Volts for 1.25 hours. Proteins were transferred to either NitroBind nitrocellulose (GE, EP4HY00010) or Immobilon-FL PVDF (Millipore, IPFL00010) membranes in Tris-Glycine transfer buffer (25 mM Tris, 192 mM glycine, 20% methanol pH 8.3) at 100 V for one hour. All primary antibodies are described in Key Resource Table.

For protein detection, membranes were blocked for one hour in 50% AquaBlock (Arlington Scientific, PP82P-500-QT-1582) in PBS (AquaBlock-P) and probed with specific antibodies (Key Resource Table) in AquaBlock-P with 0.1% Tween 20 (AquaBlock-PT) at 4 °C overnight. The membranes were subjected to four 5-minutes washes in PBS with 0.1% Tween 20 (PBS-T) at room temperature. Membranes were then incubated with secondary antibodies in AquaBlock-PT for one hour at room temperature, and subjected to four washes in PBS-T as described above. Antibody labeling was detected using an Azure Sapphire Biomolecular Imager.

### Immunofluorescence

Cells were allowed to attach to poly-L lysing coated glass slides for 2 hours. Cells were fixed with 4% paraformaldehyde in PBS pH 7.4 for 10 min at room temperature, then washed 3X with ice-cold PBS. Cells were Permeabilized for 10 min with PBS containing either 0.1% Triton X-100 (PBST), and washed 3X with PBS at room temperature. Prior to antibody addition, slides were incubated for 30 minutes with 10% serum from the species the secondary antibody was raised in. Slides were incubated in diluted primary antibody in 1% BSA in PBST in a humidified chamber overnight at 4°C. Slides were washed three times in PBS, 5 min each wash. Slides were incubated with secondary antibody in 1% BSA for 1 h at room temperature in the dark. Slides were wash three times with PBS for 5 min each in the dark. All primary antibodies are described in the Key Resource Table.

### CRISPR/Cas9 engineering

Using Lipofectamine 2000 (Invitrogen), according to the manufacturer’s recommendations, we co-transfected HTD114 cells with plasmids encoding GFP, sgRNAs and Cas9 endonuclease (Origene). Each plasmid encoded sgRNA was designed to bind at the indicated locations (Figure 1-figure supplement 1). 48h after transfection, cells were plated at clonal density and allowed to expand for 2-3 weeks. The presence of deletions was identified by PCR using the primers described in Figure 1-figure supplement 1. The single cell colonies that grew were analyzed for heterozygous deletions by PCR (Figure 1-figure supplement 1). We used retention of a heterozygous SNP to identify the disrupted allele, and homozygosity at the SNP confirmed that cell clones were homogenous.

### PCR analysis

Genomic DNA was isolated from tissue culture cells using TRIZOL Reagent (Invitrogen). PCR was performed in a 12.5µL volume using 50-100ng of genomic DNA, 1x Standard Taq Buffer (New England Biolabs, Inc.), 200µM each deoxynucleotide triphosphates, 0.2µM of each primer, and 3 units of Taq DNA Polymerase (New England Biolabs, Inc.) under the following reaction conditions: 95°C for 2 minutes, followed by 30-40 cycles of 95°C for 30 seconds, 55-62°C for 45 seconds, and 72°C for 1 minute, with a final extension time of 10 minutes at 72°C. PCR products were separated on 1% agarose gels, stained with ethidium bromide, and photographed under ultraviolet light illumination. Sequencing of PCR products was carried out at the Vollum Institute DNA Sequencing Core facility. All PCR primers are described in Figure 1-source data 3.

### DNA FISH

Mitotic chromosome spreads were prepared as described previously ^29^. After RNase (100µg/ml) treatment for 1h at 37^°^C, slides were washed in 2XSSC and dehydrated in an ethanol series and allowed to air dry. Chromosomal DNA on the slides was denatured at 75^°^C for 3 minutes in 70% formamide/2XSSC, followed by dehydration in an ice-cold ethanol series and allowed to air dry. BAC and Fosmid DNAs were labeled using nick translation (Vysis, Abbott Laboratories) with Spectrum Orange-dUTP, Spectrum Aqua-dUTP or Spectrum Green-dUTP (Vysis). Final probe concentrations varied from 40-60 ng/µl. Centromeric probe cocktails (Vysis) and/or whole chromosome paint probes (Metasystems) plus BAC or Fosmid DNAs were denatured at 75^°^C for 10 minutes and prehybridized at 37^°^C for 10 minutes. Probes were applied to denatured slides and incubated overnight at 37^°^C. Post-hybridization washes consisted of one 3-minute wash in 50% formamide/2XSSC at 40^°^C followed by one 2-minute rinse in PN (0.1M Na_2_HPO_4_, pH 8.0/2.5% Nonidet NP-40) buffer at RT. Coverslips were mounted with Prolong Gold antifade plus DAPI (Invitrogen) and viewed under UV fluorescence (Olympus). All BAC and Fosmid probes are described in Figure 1-source data 3.

### RNA-DNA FISH

Cells were plated on Poly-L-Lysine coated (Millipore Singa) glass microscope slides at ∼50% confluence and incubated for 4 hours in complete media in a 37°C humidified CO_2_ incubator. Slides were rinsed 1X with sterile RNase free PBS. Cell Extraction was carried out using ice cold solutions as follows: Slides were incubated for 30 seconds in CSK buffer (100mM NaCl/300mM sucrose/3mM MgCl_2_/10mM PIPES, pH 6.8), 10 minutes in CSK buffer/0.1% Triton X-100, followed by 30 seconds in CSK buffer. Cells were then fixed in 4% paraformaldehyde in PBS for 10 minutes and stored in 70% EtOH at -20°C until use. Just prior to RNA FISH, slides were dehydrated through an EtOH series and allowed to air dry. Denatured probes were prehybridized at 37°C for 10 min, applied to non-denatured slides and hybridized at 37°C for 14-16 hours. Post-hybridization washes consisted of one 3-minute wash in 50% formamide/2XSSC at 40^°^C followed by one 2-minute rinse in 2XSSC/0.1% TX-100 for 1 minute at RT. Slides were then fixed in 4% paraformaldehyde in PBS for 5 minutes at RT, and briefly rinsed in 2XSSC/0.1% TX-100 at RT. Coverslips were mounted with Prolong Gold antifade plus DAPI (Invitrogen) and slides were viewed under UV fluorescence (Olympus). Z-stack images were generated using a Cytovision workstation. After capturing RNA FISH signals, the coverslips were removed, the slides were dehydrated in an ethanol series, and then processed for DNA FISH, beginning with the RNase treatment step, as described above. All BAC and Fosmid probes are described in Figure 1-source data 3.

### Chromosome replication timing assay

The BrdU replication timing assay was performed as described previously on exponentially dividing cultures and asynchronously growing cells ^30^. Mitotic chromosome spreads were prepared and DNA FISH was performed as described above. The incorporated BrdU was then detected using a FITC-labeled anti-BrdU antibody (Roche). Coverslips were mounted with Prolong Gold antifade plus DAPI (Invitrogen), and viewed under UV fluorescence. All images were captured with an Olympus BX Fluorescent Microscope using a 100X objective, automatic filter-wheel and Cytovision workstation. Individual chromosomes were identified with either chromosome-specific paints, centromeric probes, BACs or by inverted DAPI staining. Utilizing the Cytovision workstation, each chromosome was isolated from the metaphase spread and a line drawn along the middle of the entire length of the chromosome. The Cytovision software was used to calculate the pixel area and intensity along each chromosome for each fluorochrome occupied by the DAPI and BrdU (FITC) signals. The total amount of fluorescent signal in each chromosome was calculated by multiplying the average pixel intensity by the area occupied by those pixels. The BrdU incorporation into human chromosomes containing CRISPR/Cas9 modifications was calculated by dividing the total incorporation into the chromosome with the deleted chromosome divided by the BrdU incorporation into the non-deleted chromosome within the same cell. Boxplots were generated from data collected from 8-12 cells per clone or treatment group. Differences in measurements were tested across categorical groupings by using the Kruskal-Wallis test ^43^ and listed as P-values for the corresponding plots. All BAC and Fosmid probes are described in Figure 1-source data 3.

### Quantification and statistical analysis

P-values were generated for eCLiP data using the “region-based” method described in^26^. For 10kb sliding windows across the genome the log2-Ratio was calculated between the number of reads in the eCLiP sample and matched control, and P-values were generated using the Python scipy.stats function “zscore”. FDR correction was performed using the Python Statsmodels function “fdrcorrection”.

---

**Figure 1-figure supplement 1. CRISPR/Cas9 deletion of the ∼7kb RBPD.** A) Integrative Genomics Viewer view showing the genomic location of ASAR6-141. The genomic locations of RNAseq and eCLIP reads from HepG2 and K562 (from ENCODE) are shown. The zoomed in view shows the ∼7kb RBPD with the location where all of the peaks of eCLIP reads map within the region (see Figure 1-source data 1). The location of sgRNAs that flank the ∼7kb RBPD are shown. The location of PCR primers (1-6) and the heterozygous SNP rs17070386 are also shown (see Figure 1-source data 3). B) Sanger sequencing traces on PCR products generated from genomic DNA using primers 5 and 6 (see Figure 1-source data 3). In previous studies we generated mouse monochromosomal hybrid cells containing the two chromosome 6 homologs from HTD114 ^7^, and sequencing traces from these two hybrids, contain either chromosome 6A (CHR6A) or chromosome 6B (CHR6B) are shown below the HTD114 panel. Sequencing traces from PCR products generated from 4 heterozygous deletions, two with CHR6A deletions {D6A(7-82-29) and D6A(7-82-13)} and two with CHR6B deletions {D6B(7-18-1) and D6B(7-63-1)} are shown. C) Junction PCR (primers 1 and 4) on genomic DNA isolated from heterozygous {D6A(7-82-29), D6A(7-82-13), D6B(7-18-1) and D6B(7-63-1)} or homozygous deletions {D6B/6A-1 and D6B/6A-2} are shown. Parental HTD114 DNA was used as control. D) Internal PCR products (primers 5 and 6) on genomic DNA isolated from heterozygous {D6A(7-82-29), D6A(7-82-13), D6B(7-18-1) and D6B(7-63-1)} or homozygous deletions {D6B/6A-1 and D6B/6A-2} are shown. Parental HTD114 DNA was used as control.

**Figure 2-figure supplement 1. Identification of mouse chromosomes containing the ∼7kb RBPD transgenes.** A) DNA FISH on mouse chromosomes from clone ∼7kb(+)A5. Mitotic spread showing DNA FISH using the ∼7kb RBPD (magenta; arrow) and a mouse chromosome 5 BAC (RP24-107D17; red; arrowheads). B and C) DNA FISH on mouse chromosomes from clone ∼7kb(+)X1. Mitotic spread showing DNA FISH using the ∼7kb RBPD (magenta; arrows) and a mouse X chromosome paint (red).

**Figure 3-figure supplement 1. RBP protein levels in cells expressing shRNAs.** K562 or HTD114 cells were transfected with empty vector (-) or shRNA against RBPs and cell extracts were processed for Western Blots using antibodies (Key Resource Table) as indicated: A and B) HNRNPU (green); C) UCHL5 (green); D and E) HNRNPC (green); F and I) HNRNPL (green); G and H) HNRNPM (green); J and K) KHSRP (green); L) HNRNPUL1 (green); M) HLTF (green); N) SAFB2 (green); O) PTBP1 (red); or P) HNRNPA1 (green). Tubulin (red) was used as loading control, and size markers are indicated.

**Figure 3-figure supplement 2. RNA FISH on cells transfected with shRNAs that did not disrupt ASAR localization.** A-D) K562 cells were transfected with empty vector (A) or shRNA expression vectors for MATR3 (B), PTBP1 (C), or PTBP2 (D). Cells were processed for RNA FISH using probes for ASAR6-141 (green; arrows) or XIST (red; arrows). E-J) HTD114 cells were transfected with empty vector (E) or shRNA expression vectors for SAFB (F), SAFB2 (G), SAFB plus SAFB2 (H), CTCF (assayed for only ASAR6 RNA) (I), or MATR3 (J). DNA was stained with DAPI, and scale bars are 10 *u*M.

**Figure 5-figure supplement 1. Disruption of the chromosome territory localization of ASAR1-187 and ASAR6 RNAs following shRNA depletion of RBPs.** HTD114 cells were transfected with empty vector (A) or shRNA expression vectors for HNRNPA1 (B), HNRNPC (C), HNRNPL (D), HNRNPM (E), HNRNPUL1 (F), KHSRP (G), HLTF (H), or UCHL5 (I). Cells were processed for RNA FISH using Fosmid probes (see Figure 1 source data 3) for ASAR1-187 (green, arrow) and ASAR6 (red, arrow). A) DNA FISH hybridization signals for chromosome 1 (CHR1 BAC DNA; magenta) and chromosome 6 (CHR6 BAC DNA; orange) are indicated by arrowheads. The brackets mark cytoplasmic regions that hybridized to both ASAR RNA FISH probes. DNA was stained with DAPI, and scale bars are 10 *u*M.

**Figure 6-figure supplement 1. Replication timing in cells transfected with shRNA expression vectors.** A and B) HTD114 cells were transfected with PTBP1 shRNA expression vector, exposed to BrdU for 5 hours, harvested for mitotic cells, and processed for BrdU incorporation (green) and DNA FISH using a chromosome 1 (A, red) or 6 (B, magenta) centromeric probe. Cells were exposed to BrdU for five hours, processed for BrdU incorporation using an antibody against BrdU. The ratio of BrdU incorporation in chromosome 1 and chromosome 6 homologs are shown in the bottom panels of A and B, respectively. C) HTD114 cells were transfected with shRNA expression vector for HNRNPU, exposed to BrdU for 5 hours, harvested for mitotic cells, and processed for BrdU incorporation (green) and DNA FISH using centromeric probes for chromosome 1 (red) or 6 (magenta). The ratio of BrdU incorporation in chromosome homologs for 1, 6, 12, and 18 are shown in the bottom panels. DNA was stained with DAPI, and scale bars are 10 *u*M.

**Figure 6-figure supplement 2. Replication timing alterations in shRNA depleted cells.** HTD114 cells were transfected with shRNA expression vectors for HNRNPA1 (A), HNRNPC (B), HNRNPL (C), HNRNPM (D), HLTF (E), KHSRP (F), HNRNPU (G), or UCHL5 (H), exposed to BrdU for 5 hours, harvested for mitotic cells, and processed for BrdU incorporation (green) and DNA FISH using a chromosome 1 centromeric probe (A and G; red, arrows) or chromosome 1 paint (B, C, D, E, F and H; red, arrows) plus a chromosome 6 centromeric probe (A-H; magenta, arrows). The brackets mark chromosomes with DRT/DMC. DNA was stained with DAPI, and scale bars are 10 *u*M.

**Figure 1-source data 1. The location of significant peaks of eCLIP reads that map within ASAR and XIST genes.**

**Figure 1-source data 2. eCLIP reads for 120 RBPs in 10 kb windows within ASARs.**

**Figure 1-source data 3. Fosmids, BACs and Primers.**

## Source Data Files

**Figure 1-figure supplement source file data.** Figure 1<u>-figure supplement 1C</u>, the original image file of the PCR products shown in panel C. Figure 1<u>-figure supplement</u> <u>1C(labels)</u>, the original image file for panel C with labels and regions of the image used are marked by the black boxes.

<u>Figure 1-figure supplement 1D</u>, the original image file of the PCR products shown in panel D. Figure 1<u>-figure supplement 1D(labels)</u>, the original image file for panel D with labels and regions of the image used are marked by the black boxes.

**Figure 3-figure supplement 1 source file data.**

<u>Figure 3-figure supplement 1A</u>, the original image file for the western blot with HNRNPU (green) and tubulin (red) antibodies on K562 cells with and without shRNA against HNRNPU. Figure 3<u>-figure supplement 1A (labels)</u>, the same image as above showing labels for the location of HNRNPU and tubulin, and the region of the image used is highlighted in white. Molecular weight standards are shown in red.

<u>Figure 3-figure supplement 1B</u>, the original image file for the western blot with HNRNPU (green) and tubulin (red) antibodies on HTD114 cells with and without shRNA against HNRNPU. Figure 3<u>-figure supplement 1B (labels),</u> the same image as above showing labels for the location of HNRNPU and tubulin, and the region of the image used is highlighted in white. Molecular weight standards are shown in red.

<u>Figure 3-figure supplement 1C</u>, the original image file for the western blot with UCHL5 (green) and tubulin (red) antibodies on K562 cells with and without shRNA against UCHL5. Figure 3<u>-figure supplement 1C (labels)</u>, the same image as above showing labels for the location of UCHL5 and tubulin, and the region of the image used is highlighted in white. Molecular weight standards are shown in red.

<u>Figure 3-figure supplement 1D</u>, the original image file for the western blot with HNRNPC (green) and tubulin (red) antibodies on K562 cells with and without shRNA against HNRNPC. Figure 3<u>-figure supplement 1C (labels)</u>, the same image as above showing labels for the location of HNRNPC and tubulin, and the region of the image used is highlighted in white. Molecular weight standards are shown in red.

<u>Figure 3-figure supplement 1E</u>, the original image file for the western blot with HNRNPC (green) and tubulin (red) antibodies on HTD114 cells with and without shRNA against HNRNPC. Figure 3<u>-figure supplement 1E (labels),</u> the same image as above showing labels for the location of HNRNPC and tubulin, and the region of the image used is highlighted in white. Molecular weight standards are shown in red.

<u>Figure 3-figure supplement 1F</u>, the original image file for the western blot with HNRNPL (green) and tubulin (red) antibodies on K562 cells with and without shRNA against HNRNPL. Figure 3<u>-figure supplement 1F (labels)</u>, the same image as above showing labels for the location of HNRNPL and tubulin, and the region of the image used is highlighted in white. Molecular weight standards are shown in red.

<u>Figure 3-figure supplement 1G</u>, the original image file for the western blot with HNRNPM (green) and tubulin (red) antibodies on K562 cells with and without shRNA against HNRNPM. Figure 3<u>-figure supplement 1G (labels)</u>, the same image as above showing labels for the location of HNRNPM and tubulin, and the region of the image used is highlighted in white. Molecular weight standards are shown in red.

<u>Figure 3-figure supplement 1H</u>, the original image file for the western blot with HNRNPM (green) and tubulin (red) antibodies on HTD114 cells with and without shRNA against HNRNPM. Figure 3<u>-figure supplement 1H (labels),</u> the same image as above showing labels for the location of HNRNPM and tubulin, and the region of the image used is highlighted in white. Molecular weight standards are shown in red.

<u>Figure 3-figure supplement 1I</u>, the original image file for the western blot with HNRNPL (green) and tubulin (red) antibodies on HTD114 cells with and without shRNA against HNRNPL. Figure 3<u>-figure supplement 1I (labels),</u> the same image as above showing labels for the location of HNRNPL and tubulin, and the region of the image used is highlighted in white. Molecular weight standards are shown in red.

<u>Figure 3-figure supplement 1J</u>, the original image file for the western blot with KHSRP (green) and tubulin (red) antibodies on K562 cells with and without shRNA against KHSRP. Figure 3<u>-figure supplement 1J (labels)</u>, the same image as above showing labels for the location of KHSRP and tubulin, and the region of the image used is highlighted in white. Molecular weight standards are shown in red.

<u>Figure 3-figure supplement 1K</u>, the original image file for the western blot with KHSRP (green) and tubulin (red) antibodies on HTD114 cells with and without shRNA against KHSRP. Figure 3<u>-figure supplement 1K (labels),</u> the same image as above showing labels for the location of KHSRP and tubulin, and the region of the image used is highlighted in white. Molecular weight standards are shown in red.

<u>Figure 3-figure supplement 1L</u>, the original image file for the western blot with HNRNPUL1 (green) and tubulin (red) antibodies on HTD114 cells with and without shRNA against HNRNPUL1. Figure 3<u>-figure supplement 1L (labels),</u> the same image as above showing labels for the location of HNRNPUL1 and tubulin, and the region of the image used is highlighted in white. Molecular weight standards are shown in red.

<u>Figure 3-figure supplement 1M</u>, the original image file for the western blot with HLTF (green) and tubulin (red) antibodies on K562 cells with and without shRNA against HLTF. Figure 3<u>-figure supplement 1M (labels)</u>, the same image as above showing labels for the location of HLTF and tubulin, and the region of the image used is highlighted in white. Molecular weight standards are shown in red.

<u>Figure 3-figure supplement 1N</u>, the original image file for the western blot with SAFB2 (green) and tubulin (red) antibodies on K562 cells with and without shRNA against SAFB2. Figure 3<u>-figure supplement 1N (labels)</u>, the same image as above showing labels for the location of SAFB2 and tubulin, and the region of the image used is highlighted in white. Molecular weight standards are shown in red.

<u>Figure 3-figure supplement 1O</u>, the original image file for the western blot with PTBP1 (red) antibody on K562 cells with and without shRNA against PTBP1. Figure 3<u>-figure</u> <u>supplement 1O (labels)</u>, the same image as above showing labels for the location of PTBP1, and the region of the image used is highlighted in white. Molecular weight standards are shown in red.

<u>Figure 3-figure supplement 1P</u>, the original image file for the western blot with HNRNPA1 (green) and tubulin (red) antibodies on K562 cells with and without shRNA against HNRNPA1. Figure 3<u>-figure supplement 1P (labels)</u>, the same image as above showing labels for the location of HNRNPA1 and tubulin, and the region of the image used is highlighted in white. Molecular weight standards are shown in red.

## RESOURCE AVAILABILITY

Further information and requests for resources and reagents should be directed to and will be fulfilled by the lead contact, Mathew Thayer

## Materials availability statement

All data generated or analyzed during this study are included in the manuscript and supporting files.

### Acknowledgments

M.J.T. was supported by NIH NIGMS (R01GM114162 and R01GM130703).

M.B.H was supported by NIH NCI 4K00CA245677-03

