## Supplementary figures and images for "ASAR lncRNAs control DNA replication timing through interactions with multiple hnRNP/RNA binding proteins"

### Figure 1 supplement 1

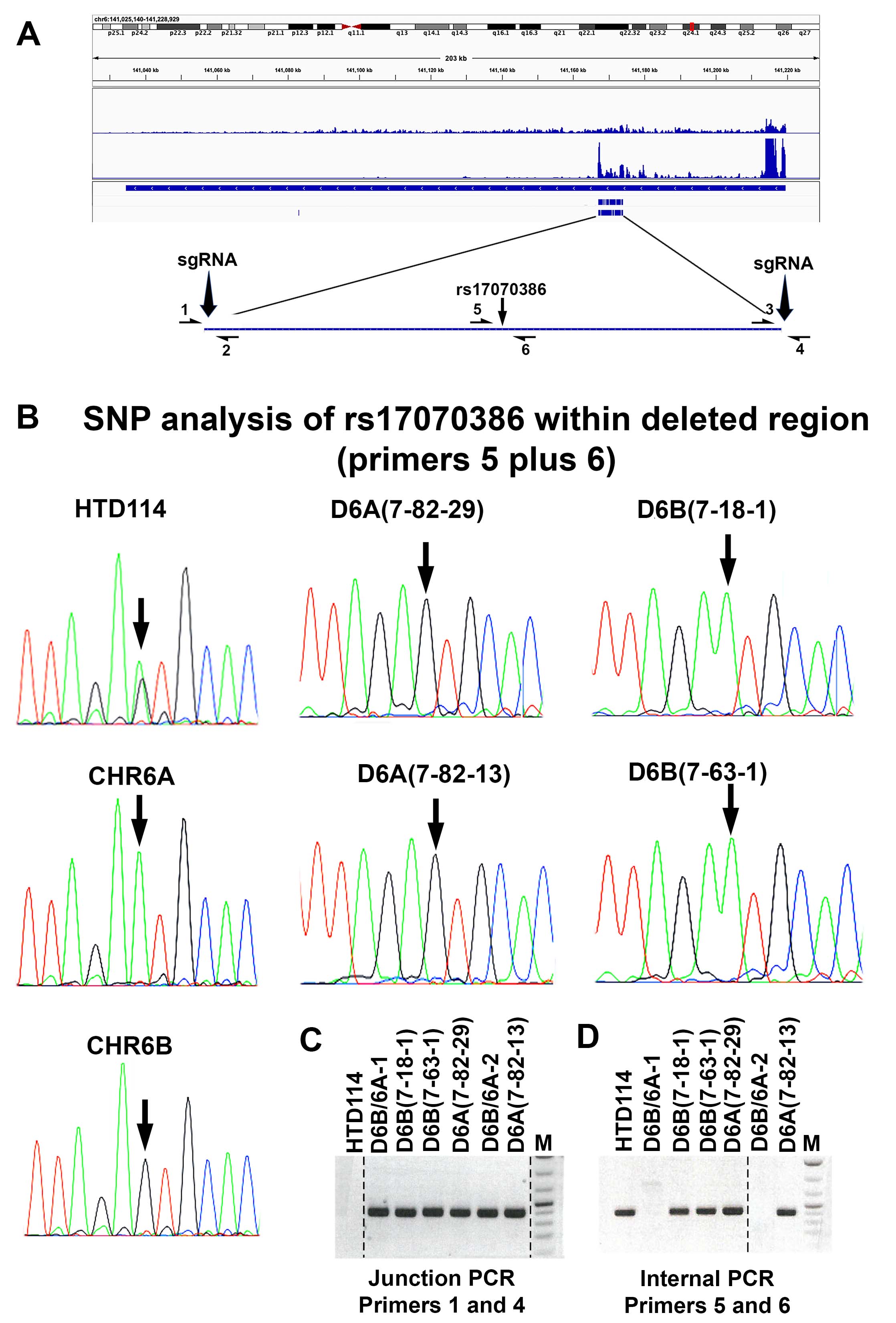

### Figure 1-figure supplement 1C copy.jpg

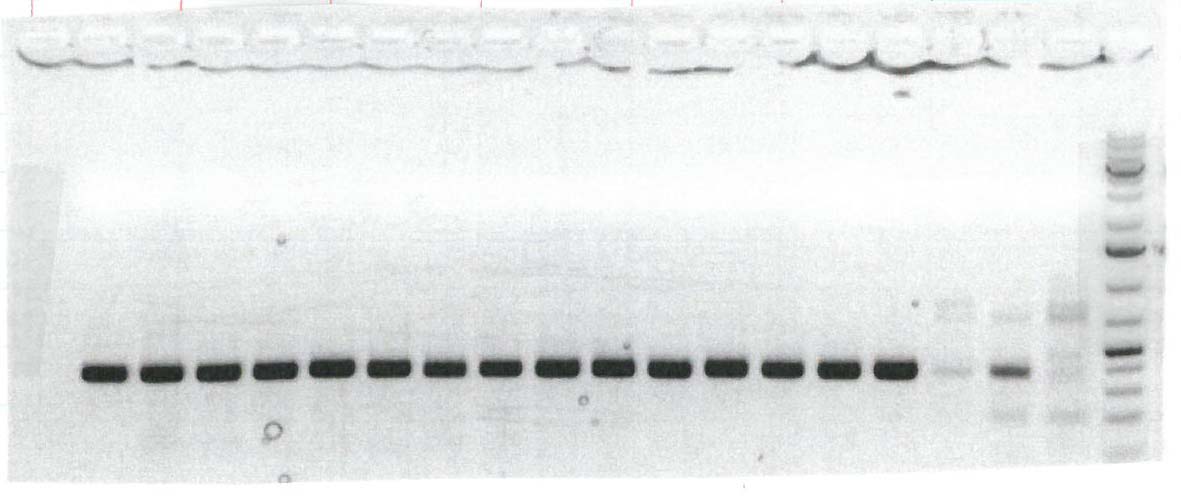

### Figure 1-figure supplement 1C(labels) copy.jpg

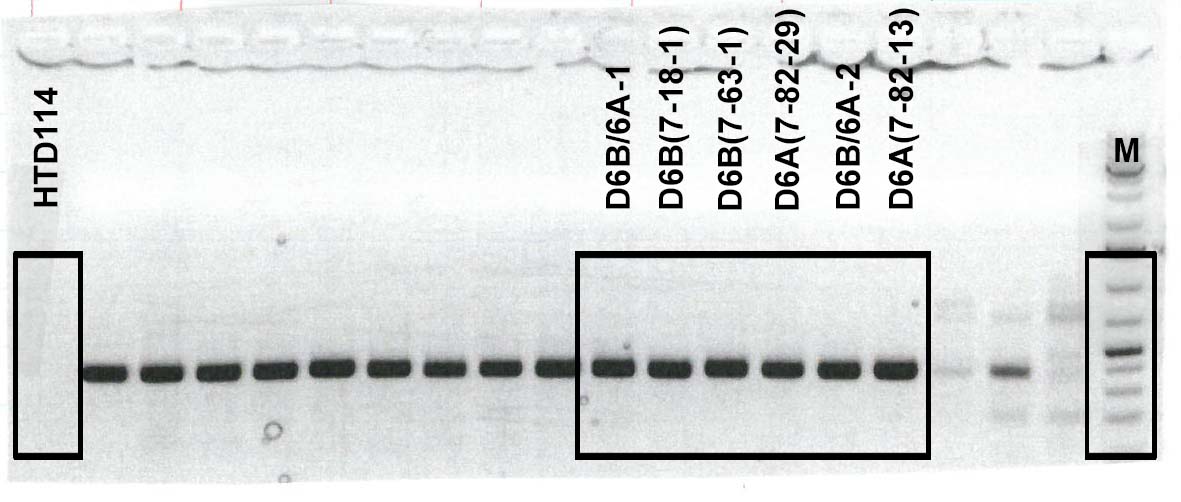

### Figure 1-figure supplement 1D copy.jpg

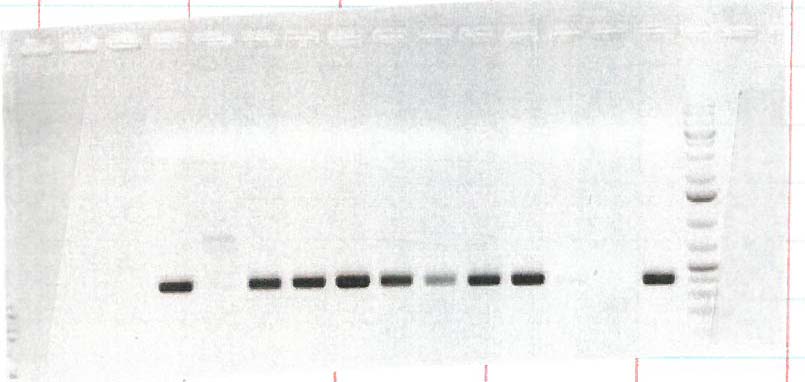

### Figure 1-figure supplement 1D(labels) copy.jpg

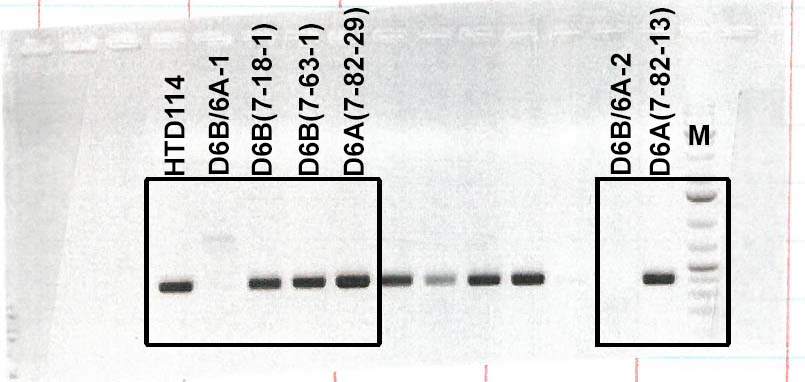

### Figure 2 supplement 1

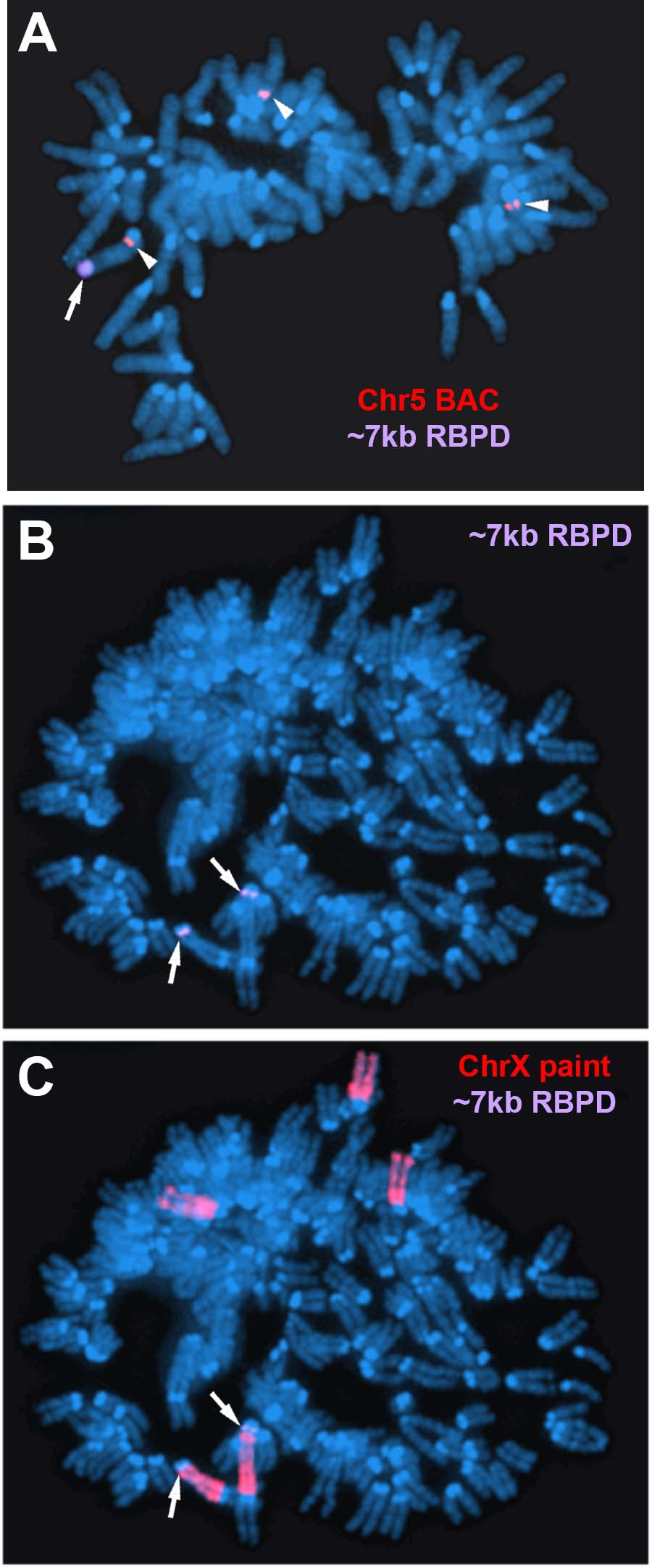

### Figure 3 supplement 1

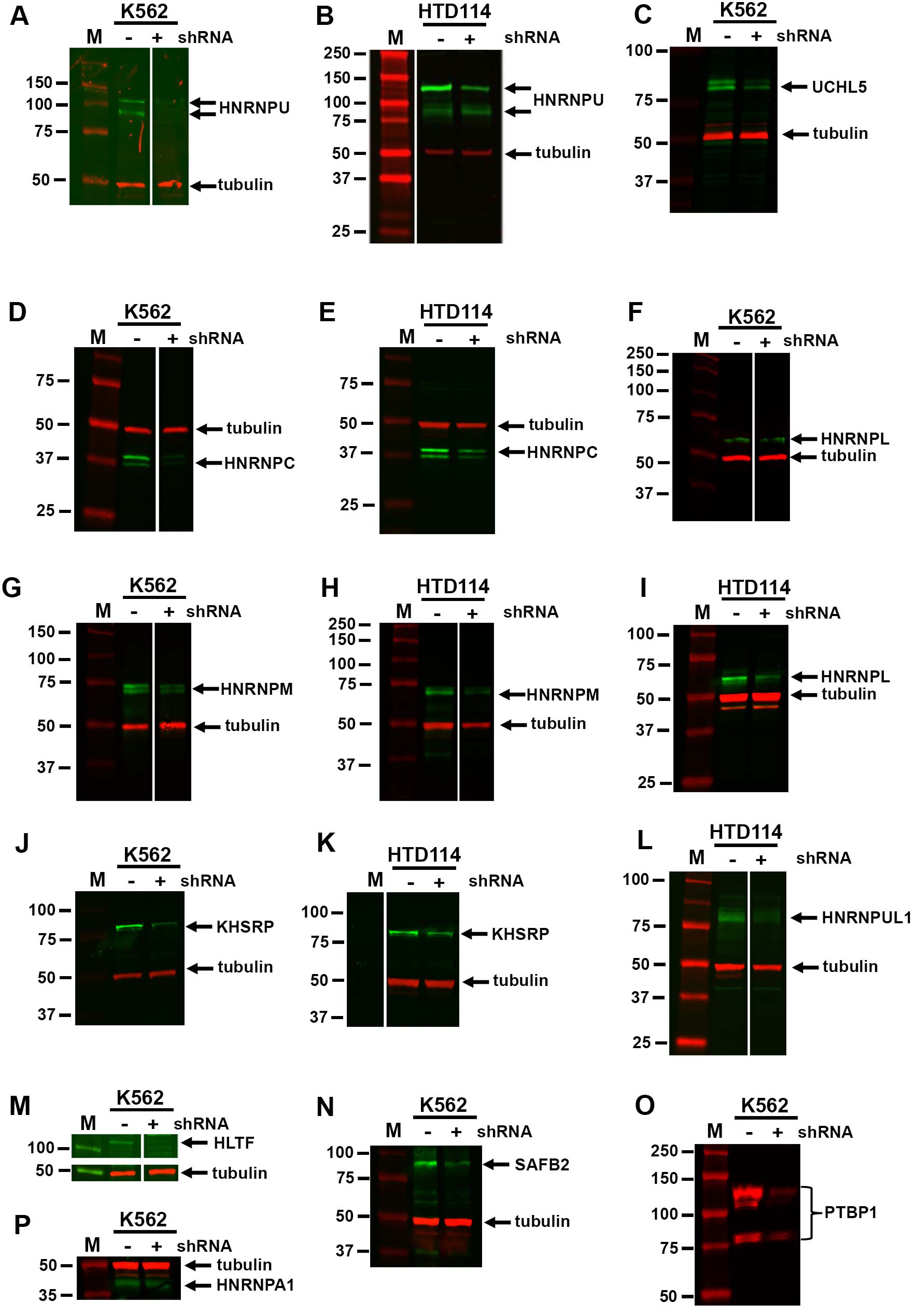

### Figure 3 supplement 2

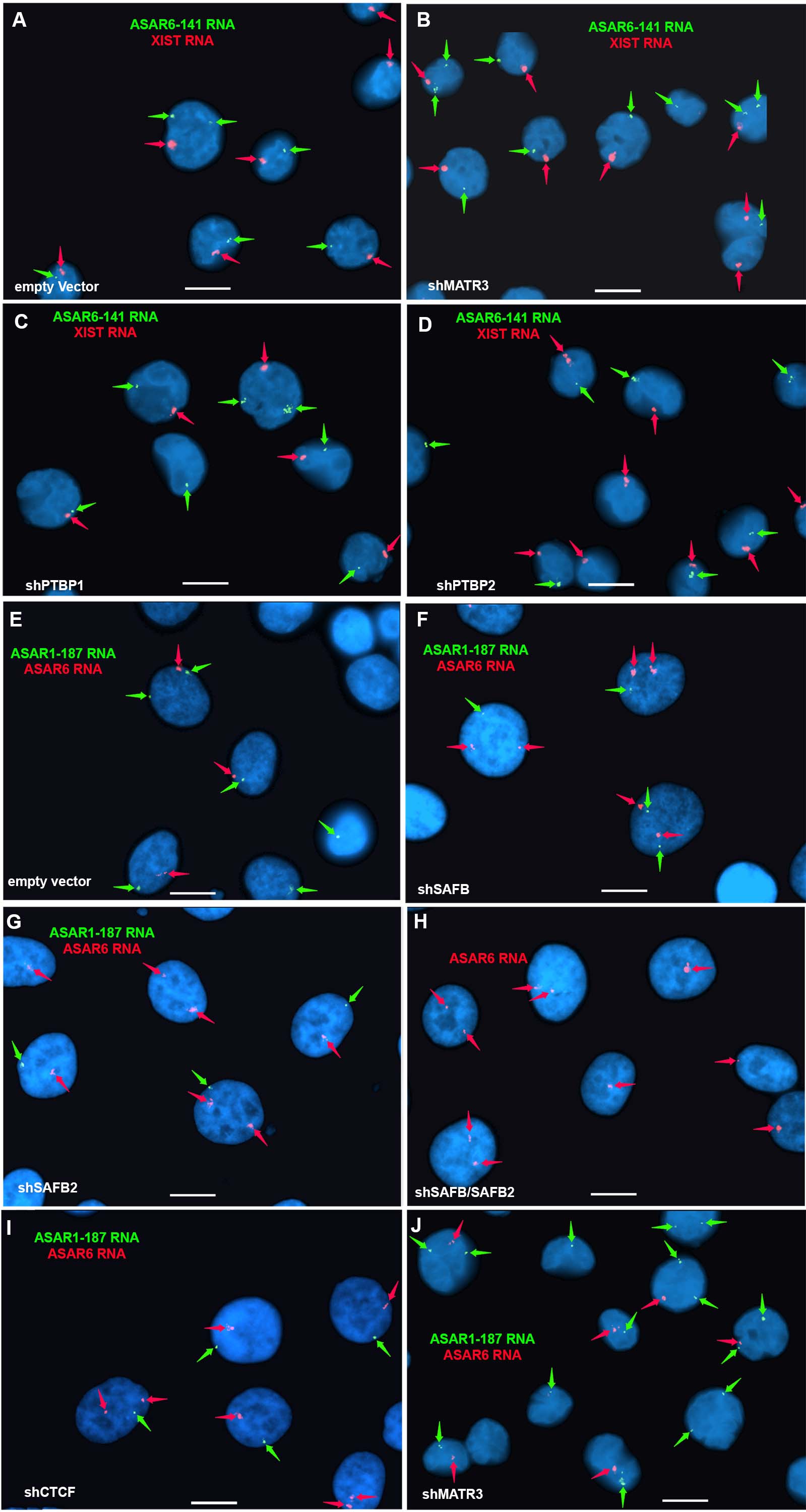

### Figure 3-figure supplement 1 A copy.jpg

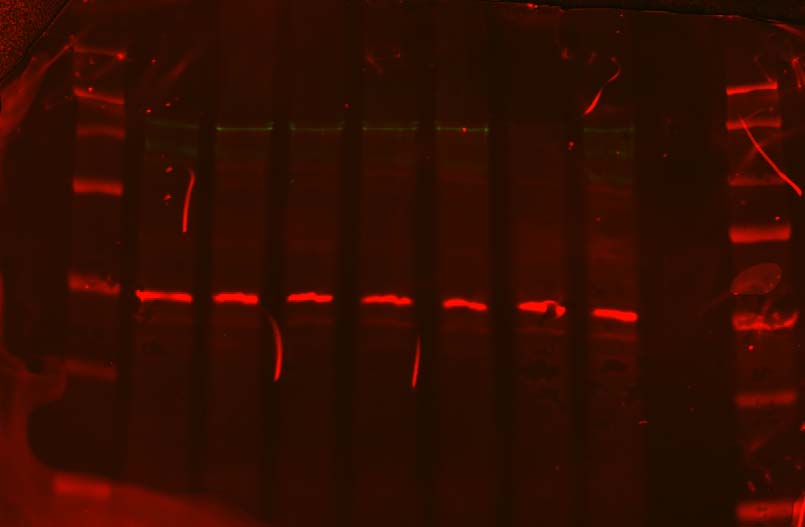

### Figure 3-figure supplement 1 A(labels) copy.jpg

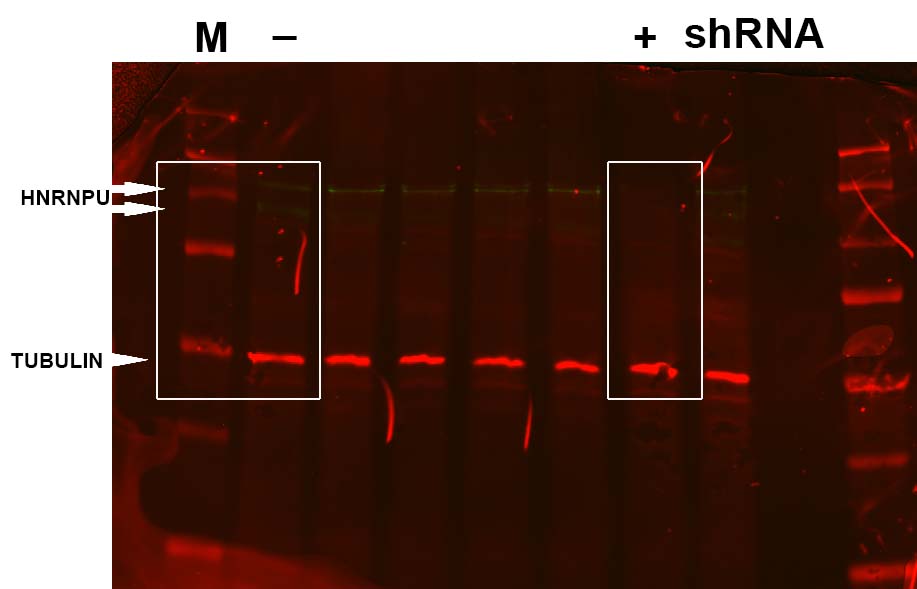

### Figure 3-figure supplement 1 B copy.jpg

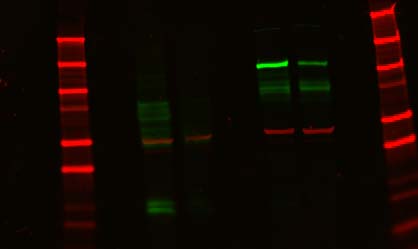

### Figure 3-figure supplement 1 B(labels) copy.jpg

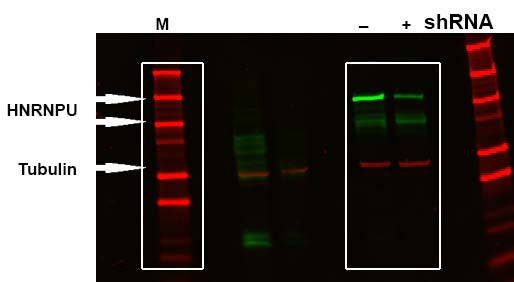

### Figure 3-figure supplement 1 C copy.jpg

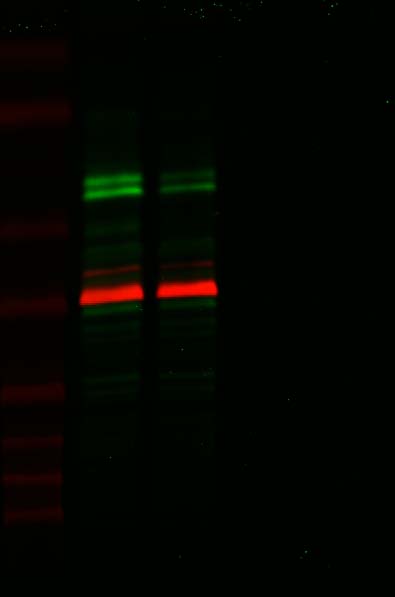

### Figure 3-figure supplement 1 C(labels) copy.jpg

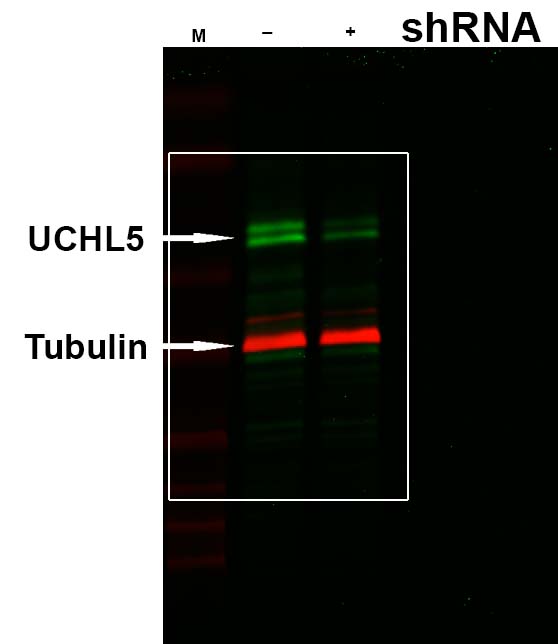

### Figure 3-figure supplement 1 D copy.jpg

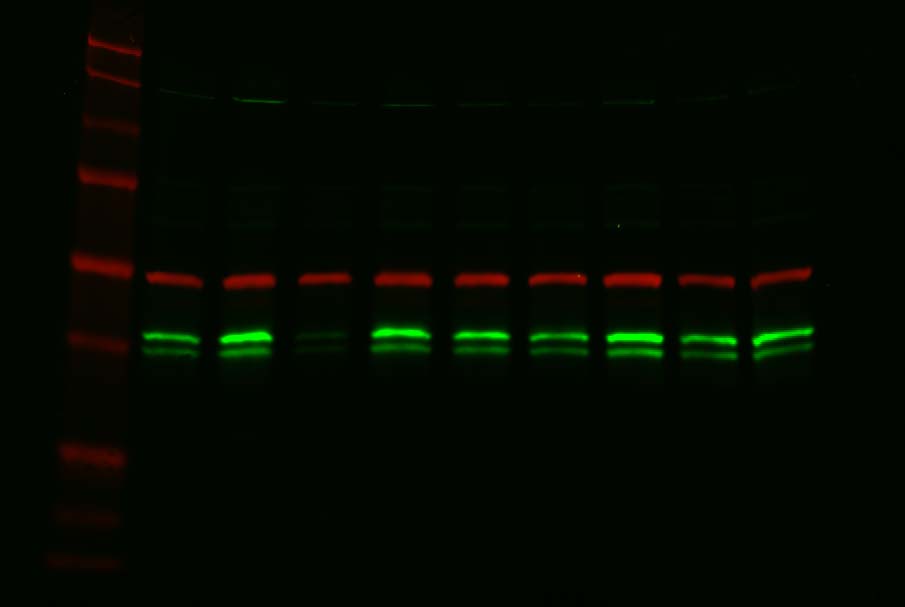

### Figure 3-figure supplement 1 D(labels) copy.jpg

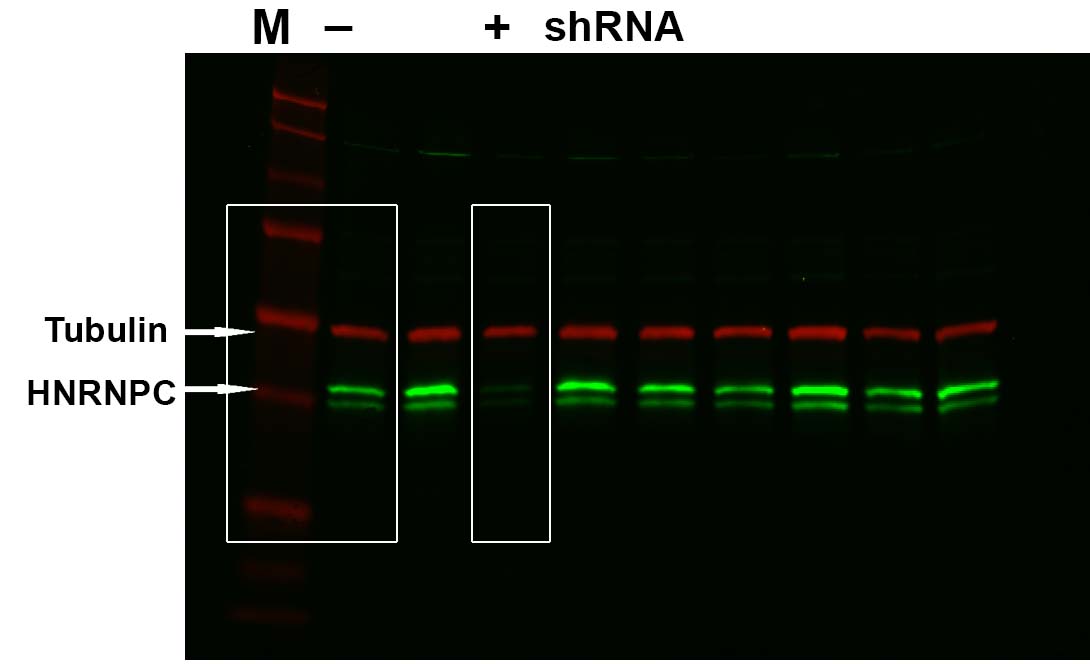

### Figure 3-figure supplement 1 E copy.jpg

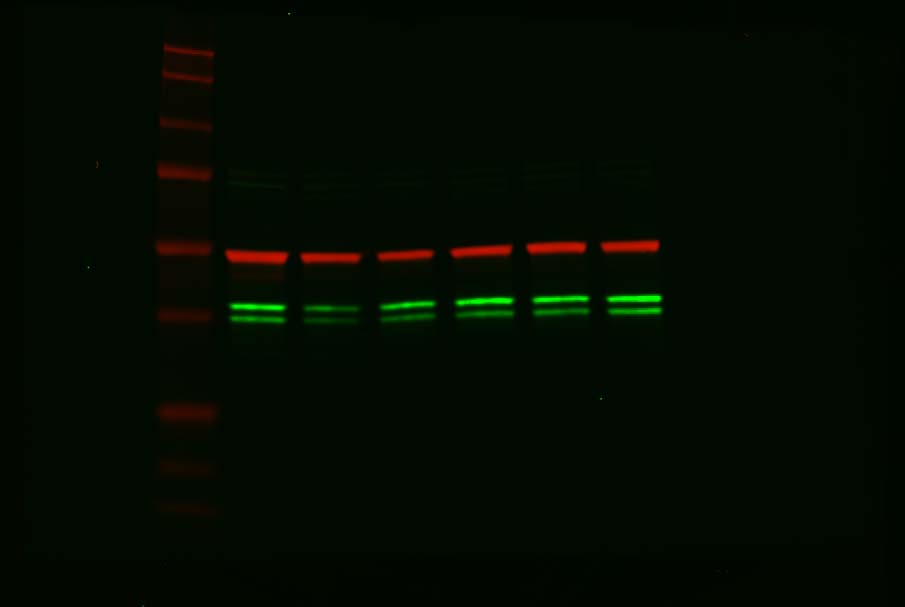

### Figure 3-figure supplement 1 E(labels) copy.jpg

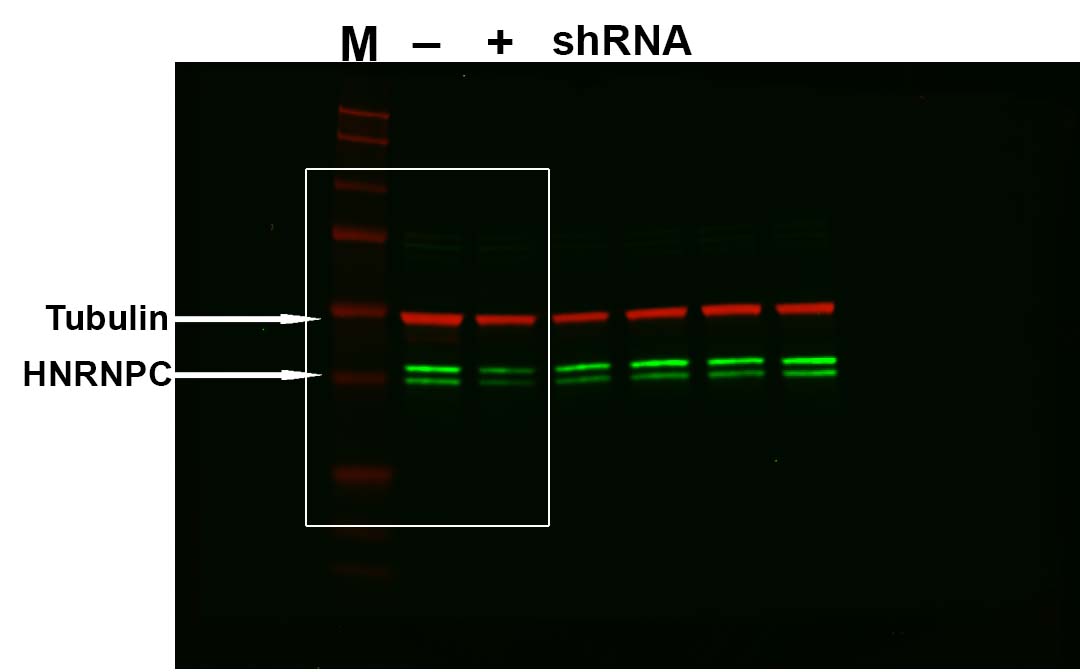

### Figure 3-figure supplement 1 F copy.jpg

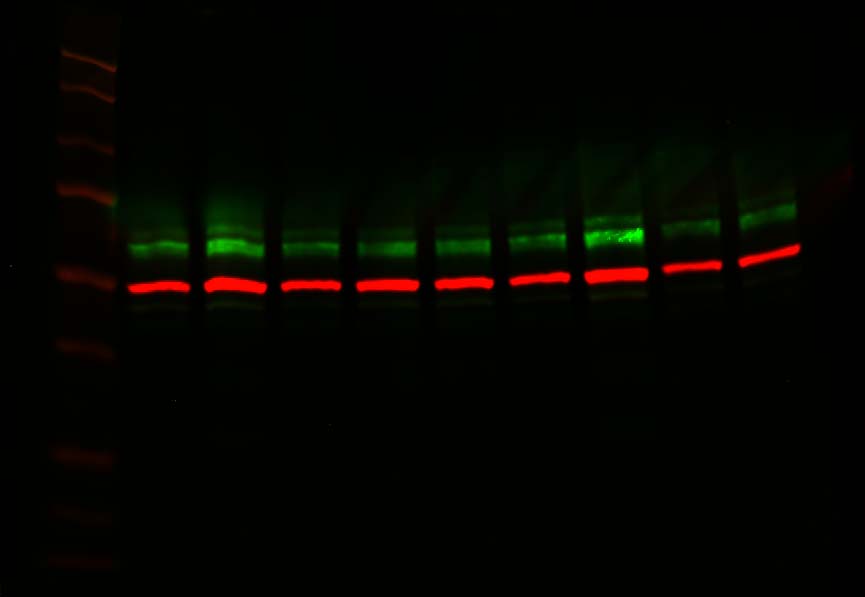

### Figure 3-figure supplement 1 F(labels) copy.jpg

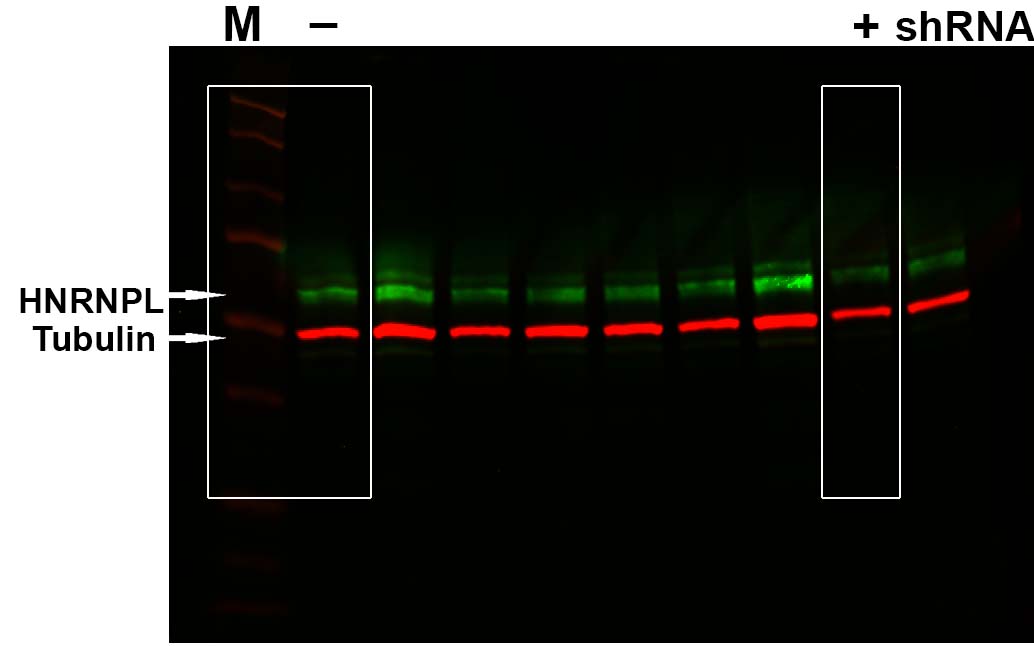

### Figure 3-figure supplement 1 G copy.jpg

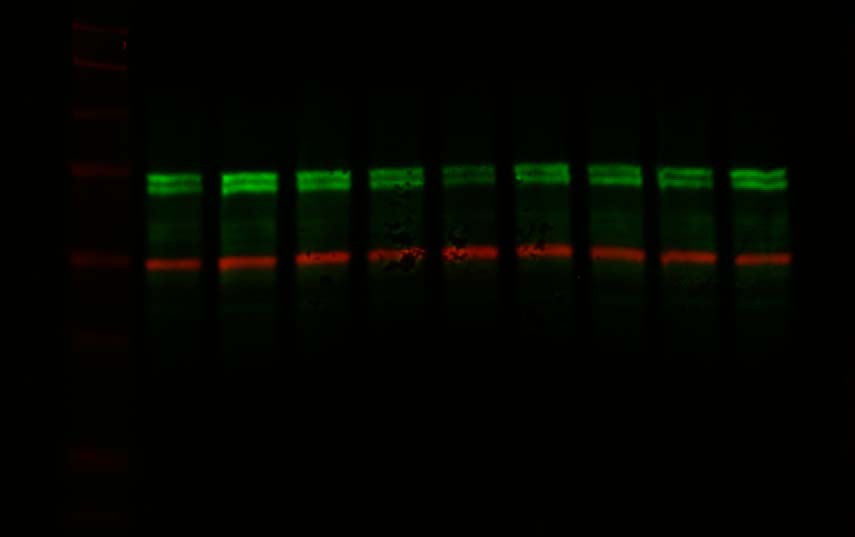

### Figure 3-figure supplement 1 G(labels) copy.jpg

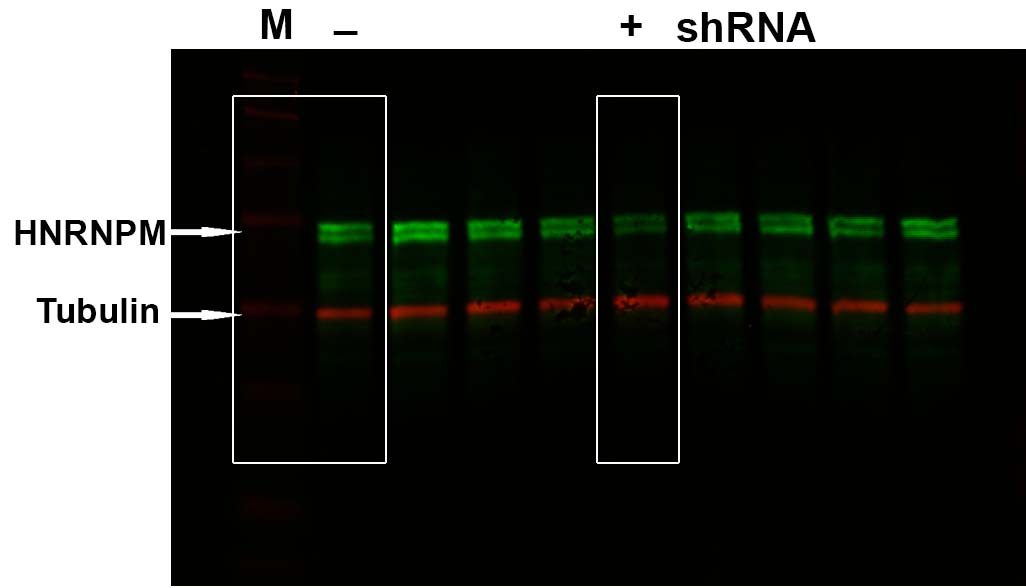

### Figure 3-figure supplement 1 H copy.jpg

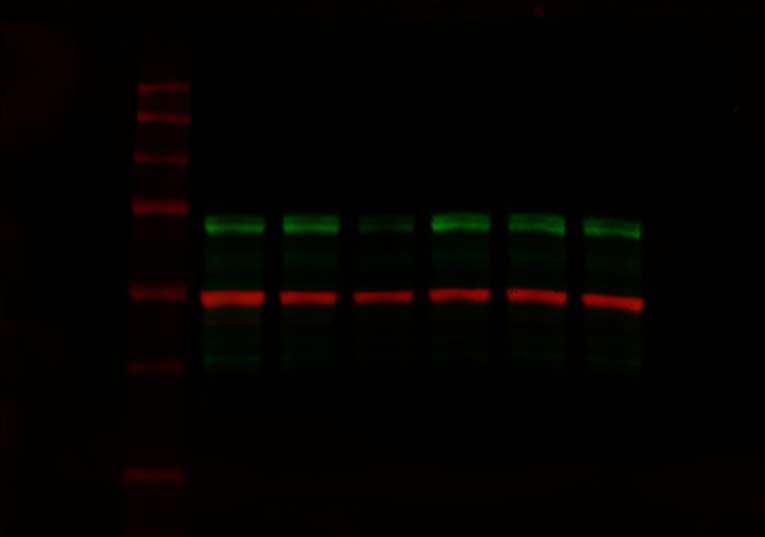

### Figure 3-figure supplement 1 H(labels) copy.jpg

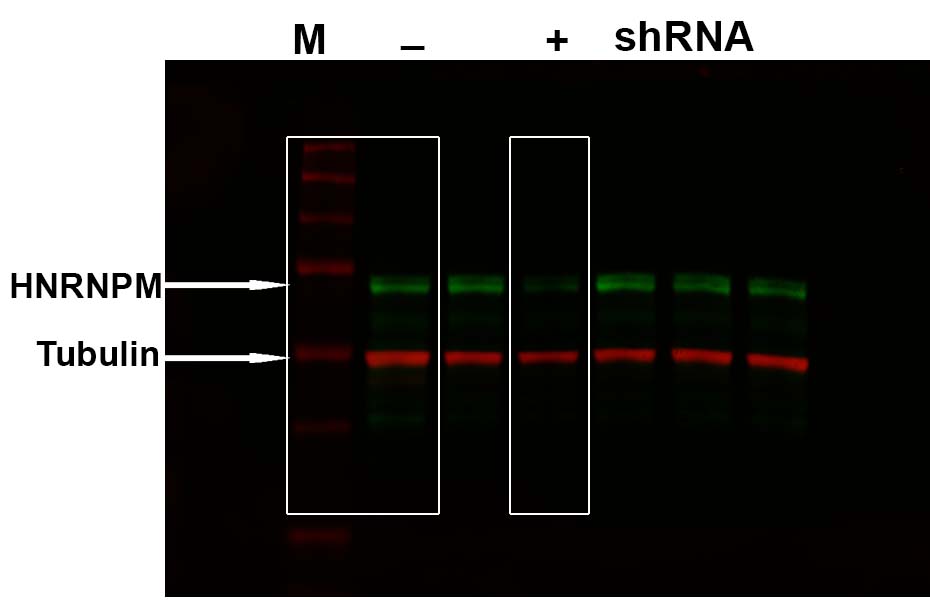

### Figure 3-figure supplement 1 I copy.jpg

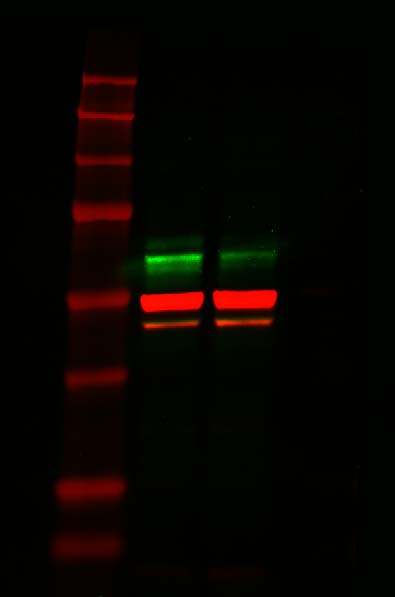

### Figure 3-figure supplement 1 I(labels) copy.jpg

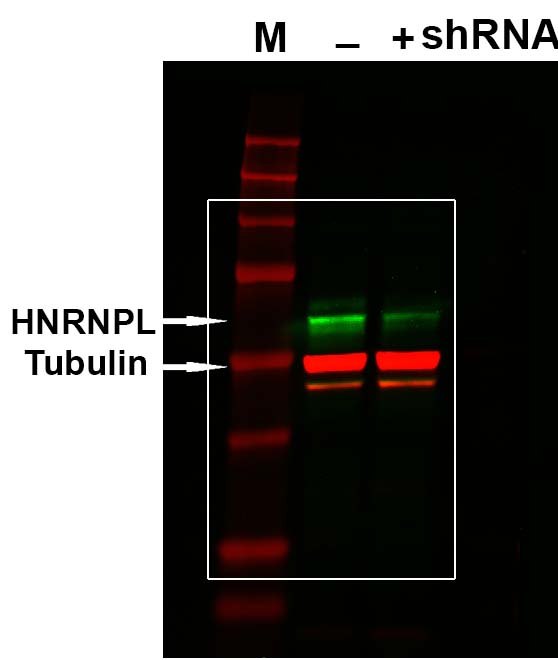

### Figure 3-figure supplement 1 J&K copy.jpg

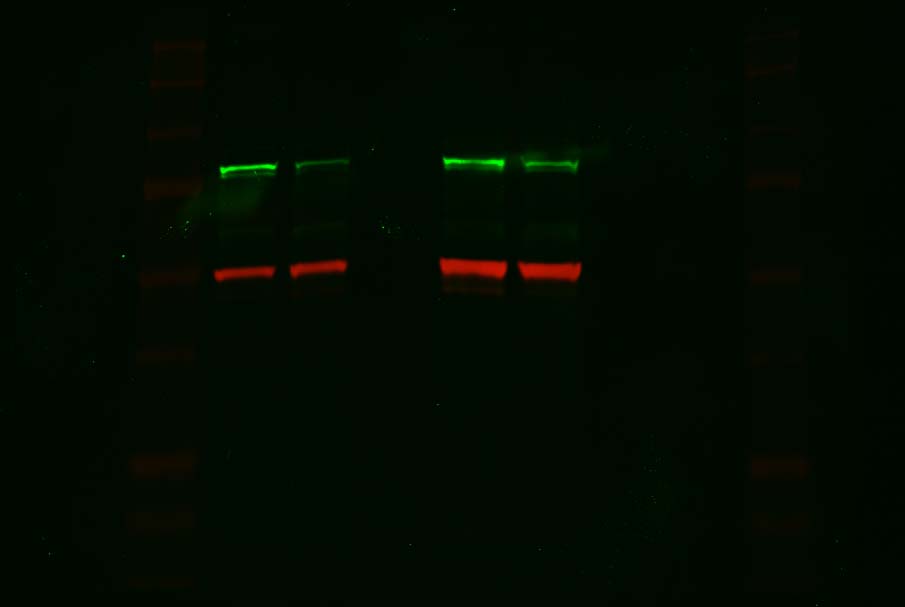

### Figure 3-figure supplement 1 J(labels) copy.jpg

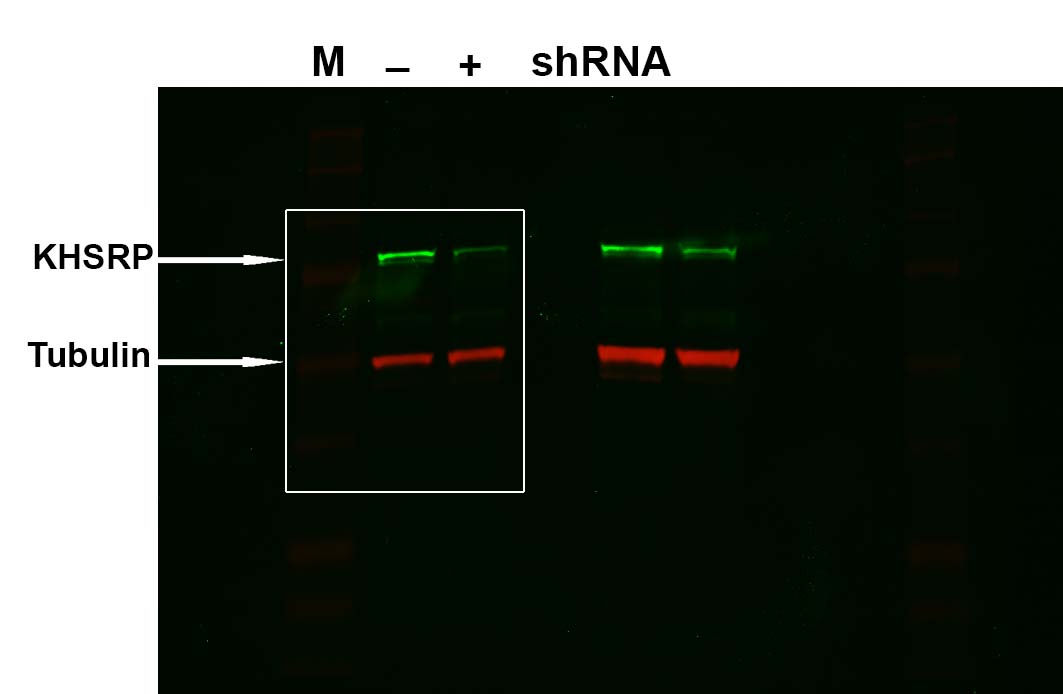

### Figure 3-figure supplement 1 K(labels) copy.jpg

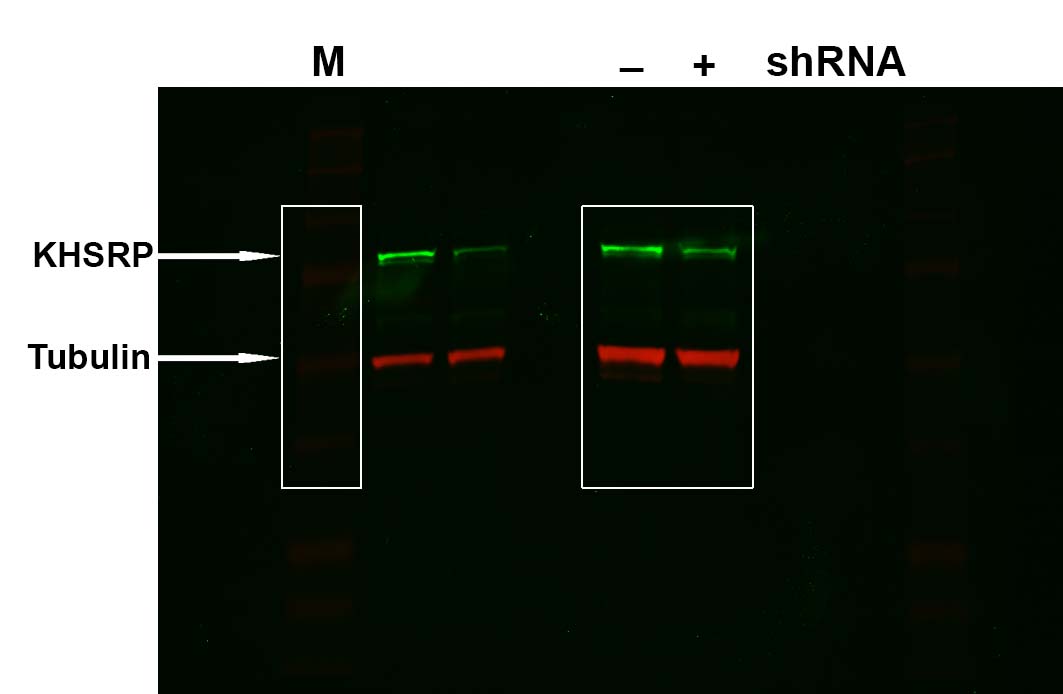

### Figure 3-figure supplement 1 L copy.jpg

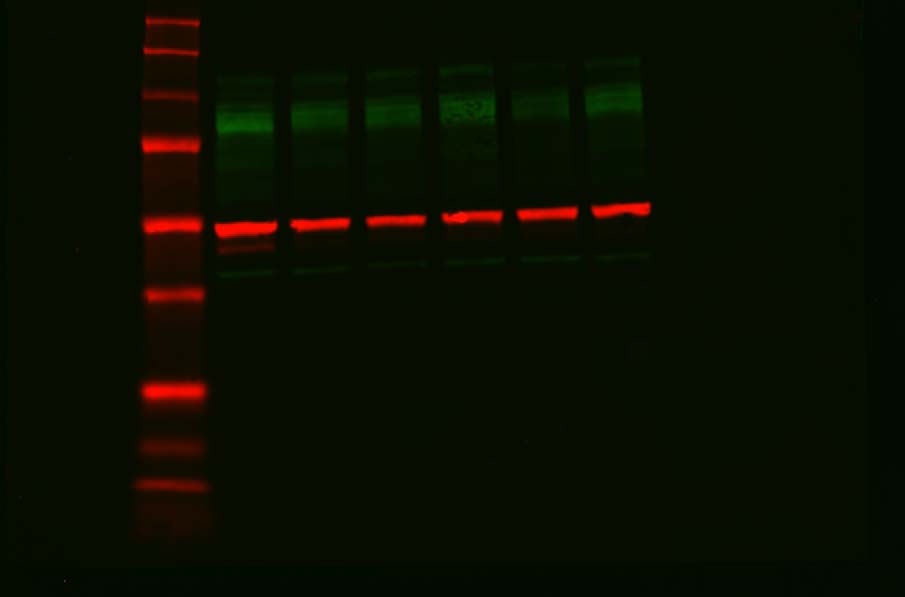

### Figure 3-figure supplement 1 L(labels) copy.jpg

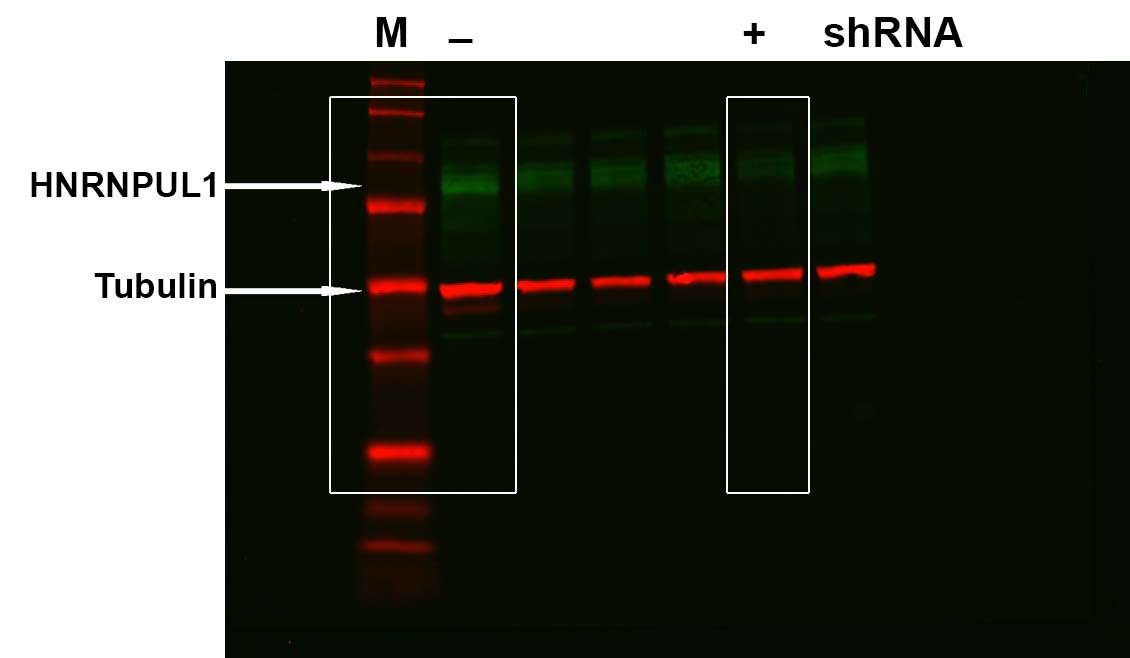

### Figure 3-figure supplement 1 M copy.jpg

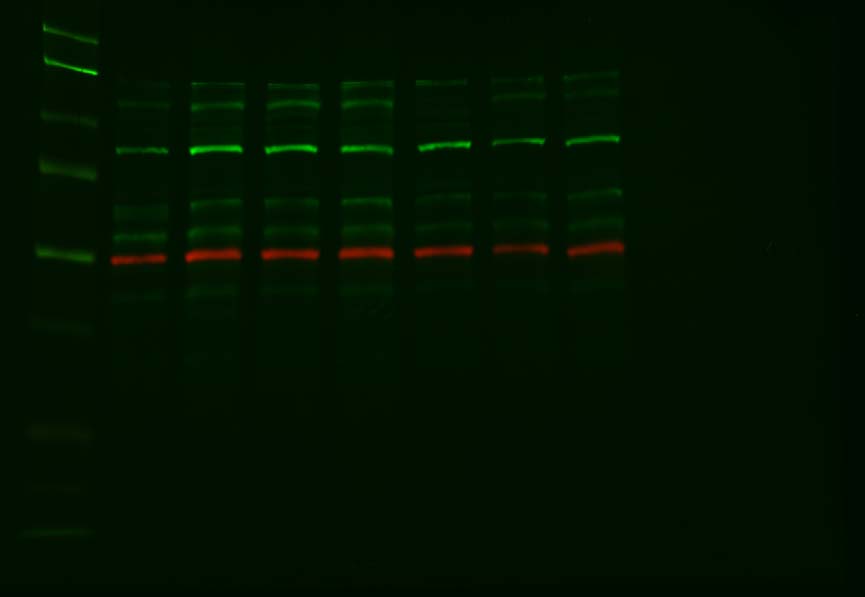

### Figure 3-figure supplement 1 M(Labels) copy.jpg

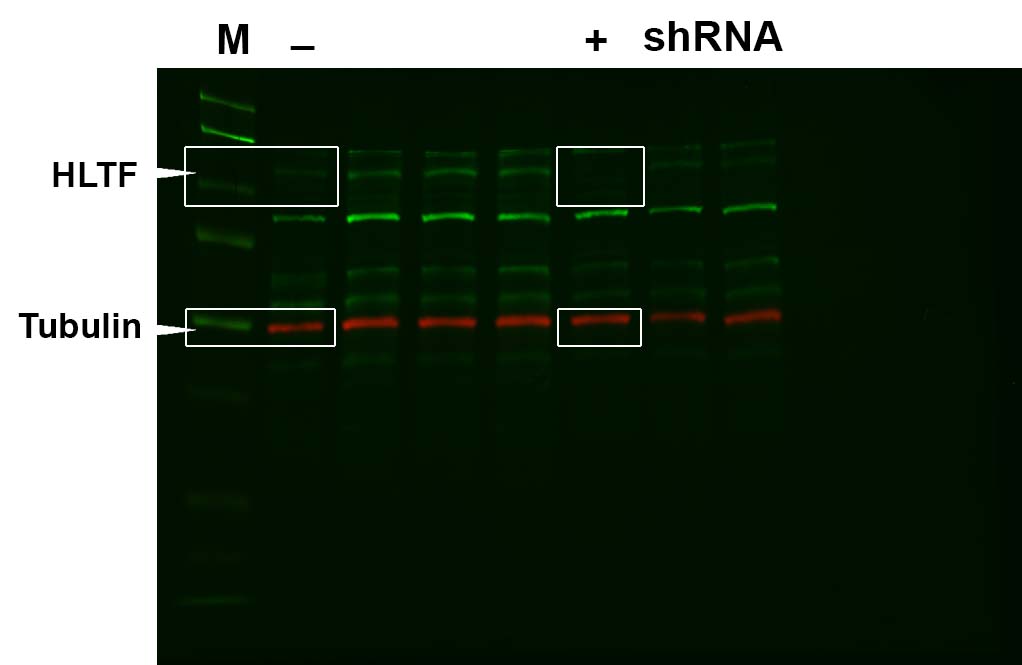

### Figure 3-figure supplement 1 N copy.jpg

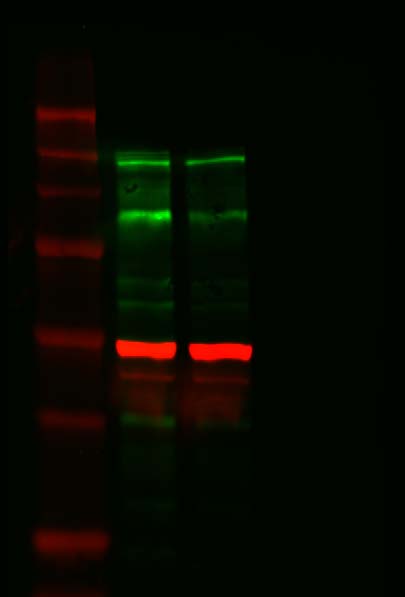

### Figure 3-figure supplement 1 N(labels) copy.jpg

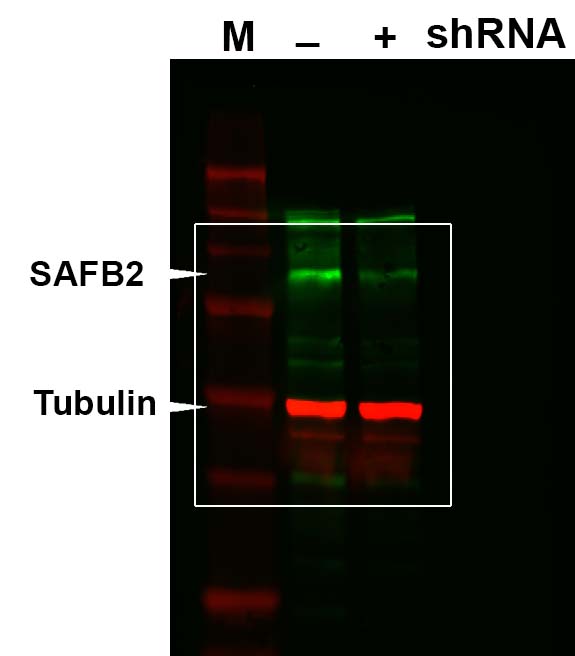

### Figure 3-figure supplement 1 O copy.jpg

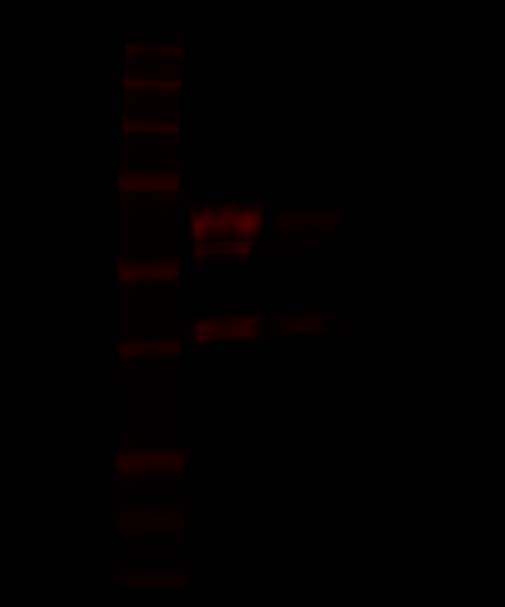

### Figure 3-figure supplement 1 O(Labels) copy.jpg

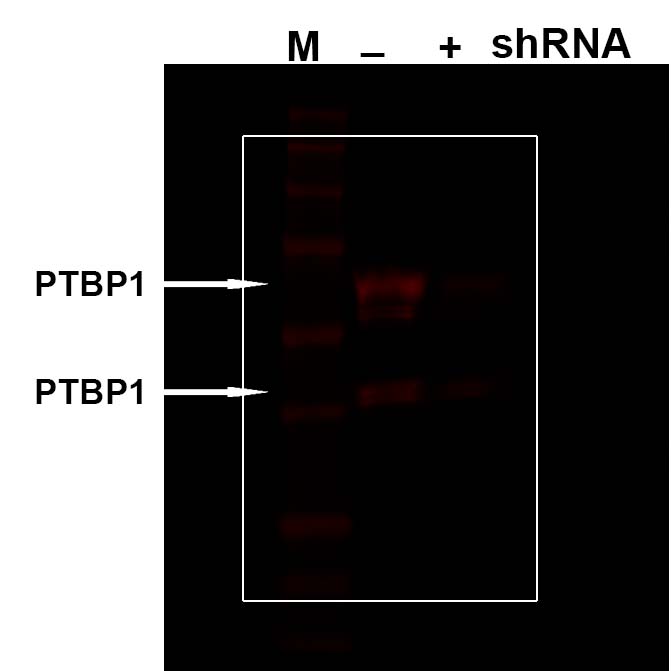

### Figure 3-figure supplement 1 P copy.jpg

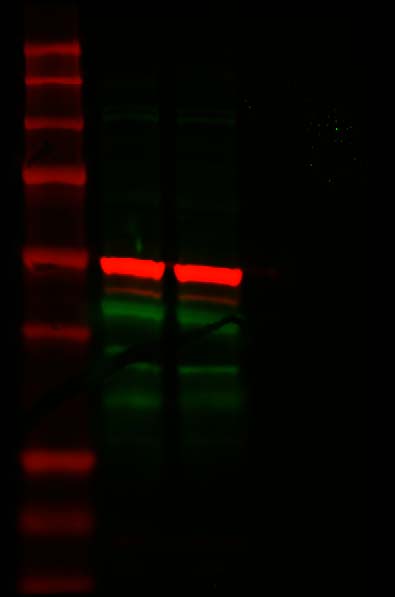

### Figure 3-figure supplement 1 P(labels) copy.jpg

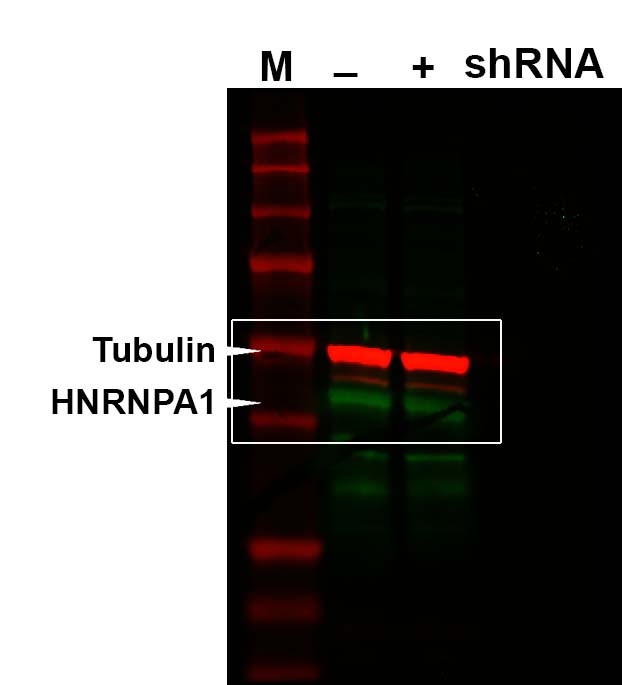

### Figure 5 supplement 1

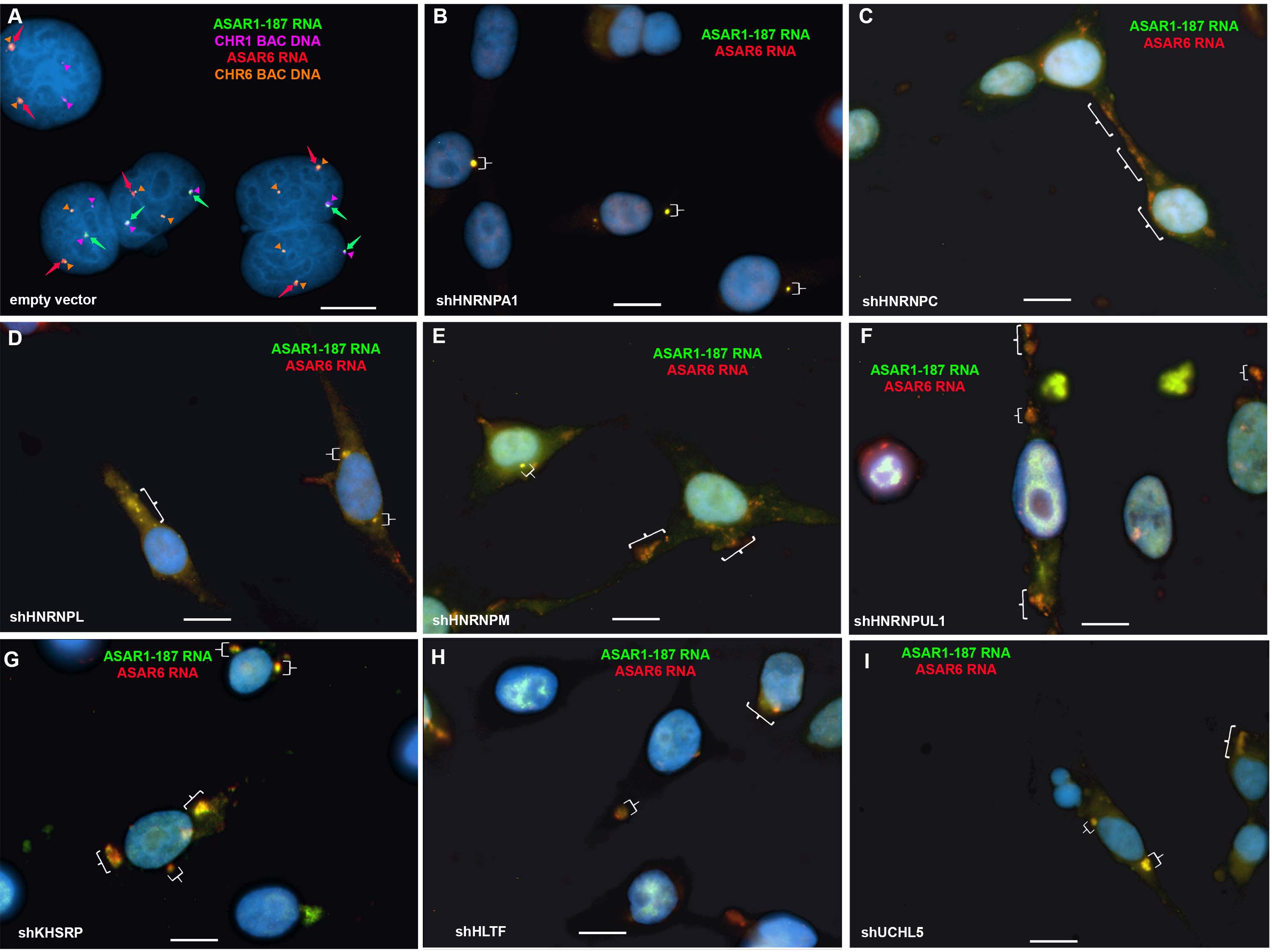

### Figure 6 supplement 1

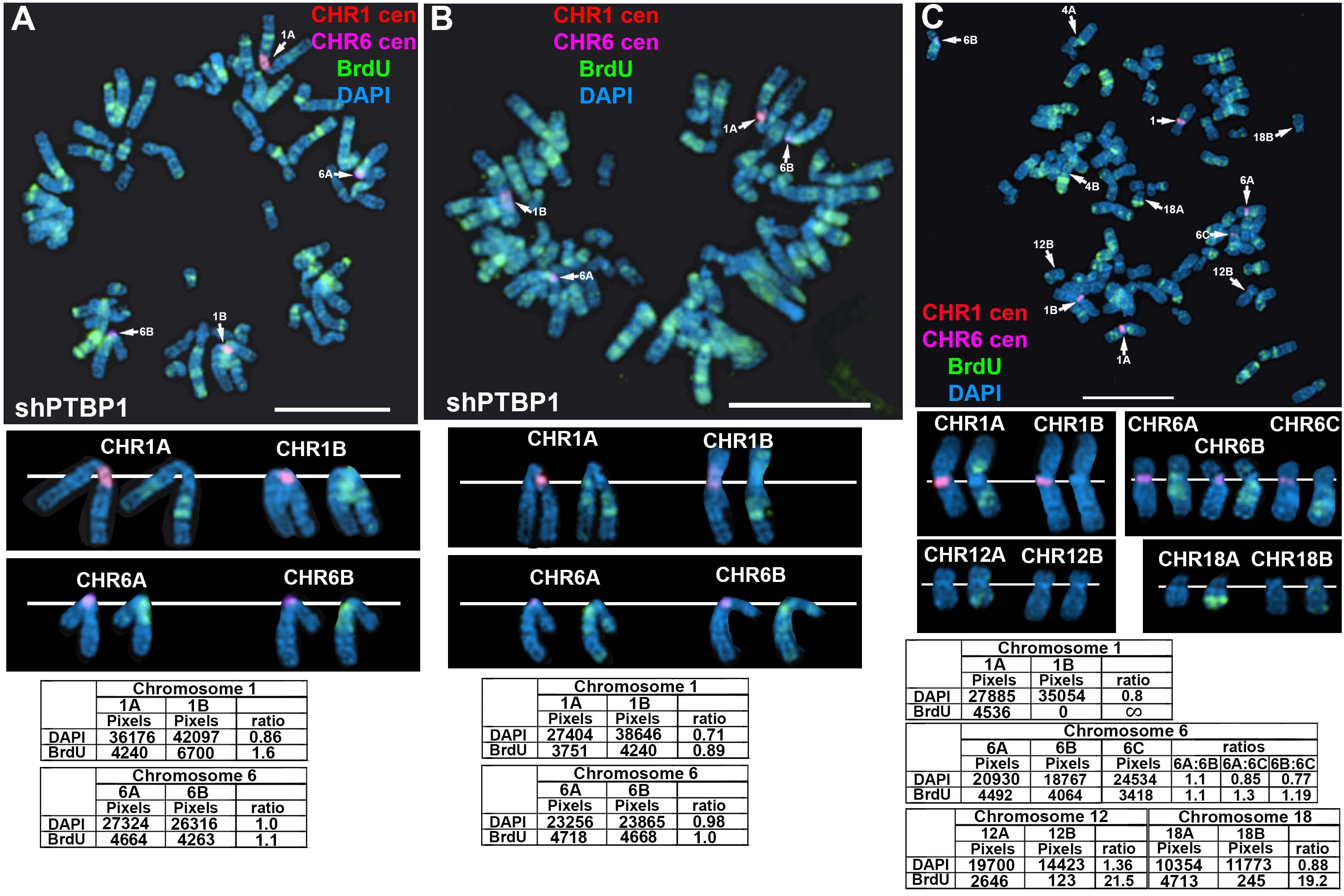

### Figure 6 supplement 2

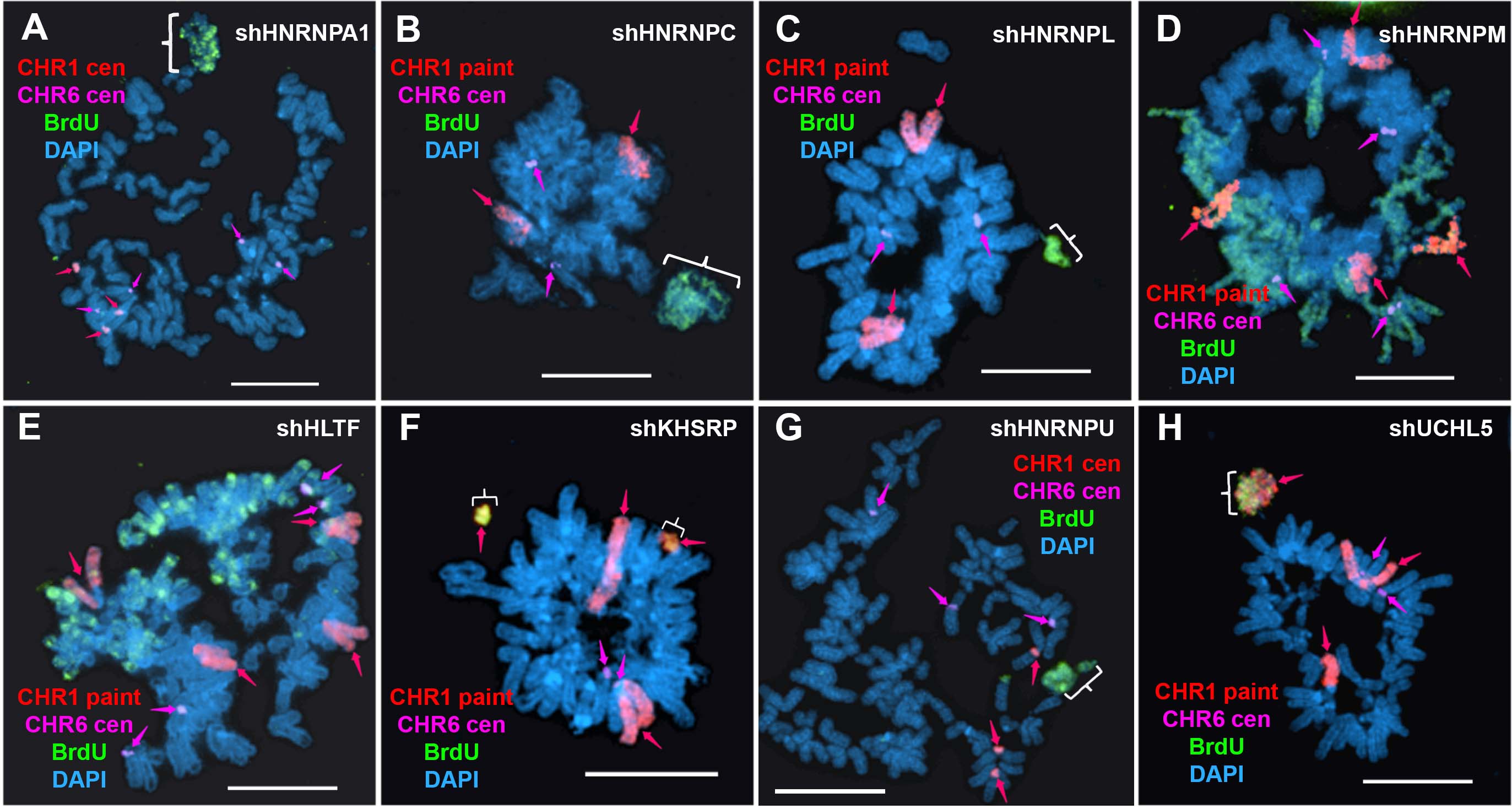
